# *MSH3* is a genetic modifier of somatic repeat instability in X-linked dystonia parkinsonism

**DOI:** 10.1101/2025.05.14.653432

**Authors:** Alan Mejia Maza, Madison Hincher, Kevin Correia, Tammy Gillis, Ayumi Nishiyama, Ellen B. Penney, Aloysius Domingo, Rachita Yadav, Micaela G. Murcar, Patrick D. Villafria Mercado, Justin S. Han, Ean P. Norenberg, Cara Fernandez-Cerado, G. Paul Legarda, Michelle Sy, Edwin Muñoz, Mark C. Ang, Cid Czarina E. Diesta, Criscely Go, Nutan Sharma, D. Cristopher Bragg, Michael E. Talkowski, Marcy E. MacDonald, Jong-Min Lee, Laurie J. Ozelius, Vanessa Chantal Wheeler

## Abstract

X-linked dystonia parkinsonism (XDP) is a progressive adult-onset neurogenerative disorder caused by the insertion of a SINE-VNTR-Alu (SVA) retrotransposon in *TAF1* gene. One element of the SVA is a tandem polymorphic CCCTCT repeat tract whose length inversely correlates with the age of disease onset. Previous observations that the repeat exhibits length-dependent somatic expansion and that XDP onset is modified by variation in DNA repair gene *MSH3* indicated that somatic repeat expansion is an important disease driver. Here, we sought to uncover genetic modifiers of CCCTCT instability in XDP patients and to provide a mechanistic link between somatic instability and disease. We determined quantitative metrics of both repeat expansion and repeat contraction in blood. Using genetic association analyses of exome sequencing data, as well as directed sequencing of a variant *MSH3* repeat, we found that *MSH3* modifies repeat expansion and contraction in blood as well as age at onset. *MSH3* alleles associated with earlier disease onset were associated with more expansion and less contraction. Conversely, alleles associated with later disease onset were associated with less expansion and more contraction. Notably, *MSH3* repeat alleles were also similarly associated with expansion and contraction in brain tissues. Our findings provide key evidence that MSH3’s role(s) in CCCTCT repeat dynamics underlies its impact on clinical disease and indicate that therapeutic strategies to lower or inhibit MSH3 are predicted to both slow CCCTCT expansion and promote CCCTCT contraction, impacting the disease course prior to clinical onset.

## Introduction

X-linked dystonia-parkinsonism (XDP) (OMIM 314250) is a fatal neurodegenerative disease endemic to the island of Panay, Philippines^1,2^. Clinically, XDP is most commonly described as a focal dystonia that presents in mid-adulthood and spreads to multiple body regions. The dystonia is accompanied, or replaced, with parkinsonism after ∼10-15 years of initial onset^3^. Studies of post-mortem XDP brains have revealed changes in the neostriatum that include the depletion of striatal medium spiny neurons (MSNs)^4–6^. MRI studies have shown changes in the cortex, substantia nigra and cerebellum^4,7–9^. Despite active research into XDP, there are no treatments to slow or prevent the disease. The genetic mutation underlying XDP is the insertion of a SINE (short interspersed nuclear element)-VNTR (variable number of tandem repeats)-Alu (SVA) retrotransposon in the TATA-box binding protein associated factor 1 (*TAF1*) gene^10,11^. The SVA insertion is associated with transcriptional dysregulation and altered splicing of *TAF1*^10,12–17^, though the causative roles of the various altered *TAF1* RNA species remain unclear. The SVA contains a polymorphic hexameric CCCTCT repeat, varying in length from 33 to 56 repeats^11,18,19^, placing XDP in a rapidly growing class of disorders associated with expanded microsatellite repeats, such as Huntington’s disease (HD)^20^. The length of the CCCTCT repeat is inversely correlated with XDP age at onset (AAO)^11,18,19^, as observed in other repeat expansion diseases, indicating a critical role of CCCTCT length in driving XDP pathogenesis. Also in common with other repeat diseases, the CCCTCT repeat is unstable in the germline and somatic tissues^11,18,19^. We recently demonstrated that somatic instability of the CCCTCT repeat is expansion-biased, tissue-specific, and repeat length-dependent, suggesting that the repeat length-dependent property driving XDP onset is its propensity to expand in somatic cells^19^.

Importantly, some individuals present AAO either earlier or later than predicted by their inherited CCCTCT length, suggesting that other genetic modifiers may also play a role in determining the rate of XDP onset. Indeed, a recent genome-wide association study (GWAS) showed that variants near or within MutS Homolog-3 *(MSH3*) and *PMS1* homolog-2 (*PMS2*) genes modify XDP AAO^21^, together accounting for ∼25% of AAO residual variance (AAO adjusted for CCCTCT repeat length). *MSH3* and *PMS2* encode proteins in the mismatch repair (MMR) pathway and were previously uncovered in GWAS, amongst other MMR/DNA repair genes, as modifiers of HD motor onset and other clinical phenotypes^22–24^. Significantly, these genes modify somatic expansion of the HD CAG repeat^23,25–27^, indicating that they modify HD by altering the rate of somatic repeat expansion in the brain. It seems likely, therefore, that *MSH3* and *PMS2* modify the somatic instability of the XDP CCCTCT repeat. However, genetic modifiers of somatic instability in XDP remain to be identified. Here, we aim to uncover genetic modifiers of CCCTCT instability in XDP patients and to provide a mechanistic link between somatic instability and disease. Combining whole exome sequencing, GWAS data and targeted genotyping, we provide direct evidence that *MSH3* modifies somatic CCCTCT expansion and contraction in XDP patients and show that the same *MSH3* variants alter both expansion, contraction and AAO, directly supporting somatic repeat dynamics as being a major driver of XDP clinical onset.

## Materials and methods

### Patients and sample collection

Patients recruited for this study included individuals with XDP evaluated at Massachusetts General Hospital (MGH) (Boston, MA, USA), Jose R. Reyes Memorial Medical Center (JRRMMC) (Manila, Philippines), and the Sunshine Care Foundation Clinic in Roxas City (Panay, Philippines). All participants provided written informed consent, and the study was approved by local Institutional Review Boards (IRBs) at both MGH and JRRMMC.

Blood samples were collected from male individuals (n = 382, of which 360 were affected and 22 were non-manifesting carriers-the latter used for modeling instability only). Genomic DNA (gDNA) was extracted from blood using the Flexigene Reagent Kit (Qiagen) according to the manufacturer’s instructions and XDP genetic status was confirmed by PCR amplification of a 48 bp deletion haplotype marker as previously described^11,19^. This study included samples used in our previous study^19^. XDP clinical diagnosis was determined by trained clinicians, as described^3^.

Post-mortem brain tissues from male individuals with XDP (n = 60) were obtained from the Collaborative Center for XDP (CCXDP) Brain Bank at MGH (Boston, MA, USA). gDNA was extracted from post-mortem brain tissues using the DNeasy Blood & Tissue Kit (Qiagen), according to the manufacturer’s instructions and including the addition of 3µl of RNase. The post-mortem brain tissues used in this study comprised up to 17 regions from up to 60 males: frontal cortex Brodmann area 9 (BA9, n = 58), caudate (n = 33), cerebellum (n = 60), cingulate gyrus (n = 20), deep cerebellar nuclei (n = 21), hippocampus (n = 19), insular cortex (n = 19), inferior olivary nucleus (n = 9), lateral thalamus (n = 20), media thalamus (n = 34), occipital cortex (n = 60), parietal cortex (n = 20), putamen (n = 19), red nucleus (n = 11), substantia nigra (n = 19), subthalamic nucleus (n = 7) and temporal pole (n = 57). A subset of these brain tissues was used for analyses of somatic expansion in our previous study^19^.

### Determination of XDP repeat length

The length of the XDP CCCTCT repeat tract in blood and brain samples was determined using a fluorescent PCR-based assay^11,19^. PCR reactions consisted of 125 ng of gDNA per reaction added in a 50 μl reaction volume with 14 μl buffer, 2 μl of 2.5 mM dNTPs, 0.5 U polymerase provided with the PrimeSTAR GLX polymerase (TAKARA) kit and 10 mM of each primer). The PCR conditions were 94°C x 2 min; 30 x (98 °C x 10 s, 64°C x 35 s). The primers used were:

Forward = 5’-[6FAM]-AGCAGTACAGTCCAGCTTTGGC-3’ and Reverse = 5’ – CTCAAGCCTTATTACAATGCCAGT – 3’.

PCR products were run on an ABI 3730xl DNA Analyzer (Applied Biosystems) using GeneScan 500 LIZ as an internal standard and output data analyzed using GeneMapper V5 (Applied Biosystems)^11,19^. The PCR products comprised a series of peaks separated by 6 bp (one CCCTCT repeat). The repeat length of the modal peak, assumed to be the inherited repeat length, was assigned based on DNA standards of known repeat lengths (37, 44, 50).

### Defining expansion and contraction phenotypes in blood and relationships to AAO

Expansion and contraction indices were obtained from fragment sizing analyses, as described previously, using a 5% relative peak height threshold cut-off^28^. We first developed regression models to explain somatic expansion or contraction indices as a function of inherited repeat length and age at sample collection and subsequently derived residual expansion and contraction index values from these models to explain the contribution of somatic expansion or contraction to AAO. We found that logarithmic transformation of the somatic expansion or contraction indices improved the model fitting.

In both the expansion and contraction modeling, we initially tested the interaction term between repeat length and age at sample collection (See **Figure S1** for plots that indicate a weak interaction). However, the coefficients obtained in models with the interaction term did not contribute substantially to the model and did not alter the outcome of downstream exome-wide association study analyses. We therefore did not include an interaction term in order to simplify the models and avoid centering the data, allowing for an easier interpretation of the residual values. For contractions, we found that inclusion of only the first contraction peak relative to the modal allele (N-1 peak) provided a better model fit (R^2^=0.4101) than models based on inclusion of either all the contraction peaks (R^2^=0.3861) or the first plus second contraction peaks (R^2^=0.3934). Thus, we used the N-1 peak inclusion model to minimize outlier effects in the downstream genetic analyses but note that all three contraction models resulted in the identification of a signal at the *MSH3* locus in the exome association analysis.

The final models used (see **Figure 1d**) were as follows:

**Figure 1.**
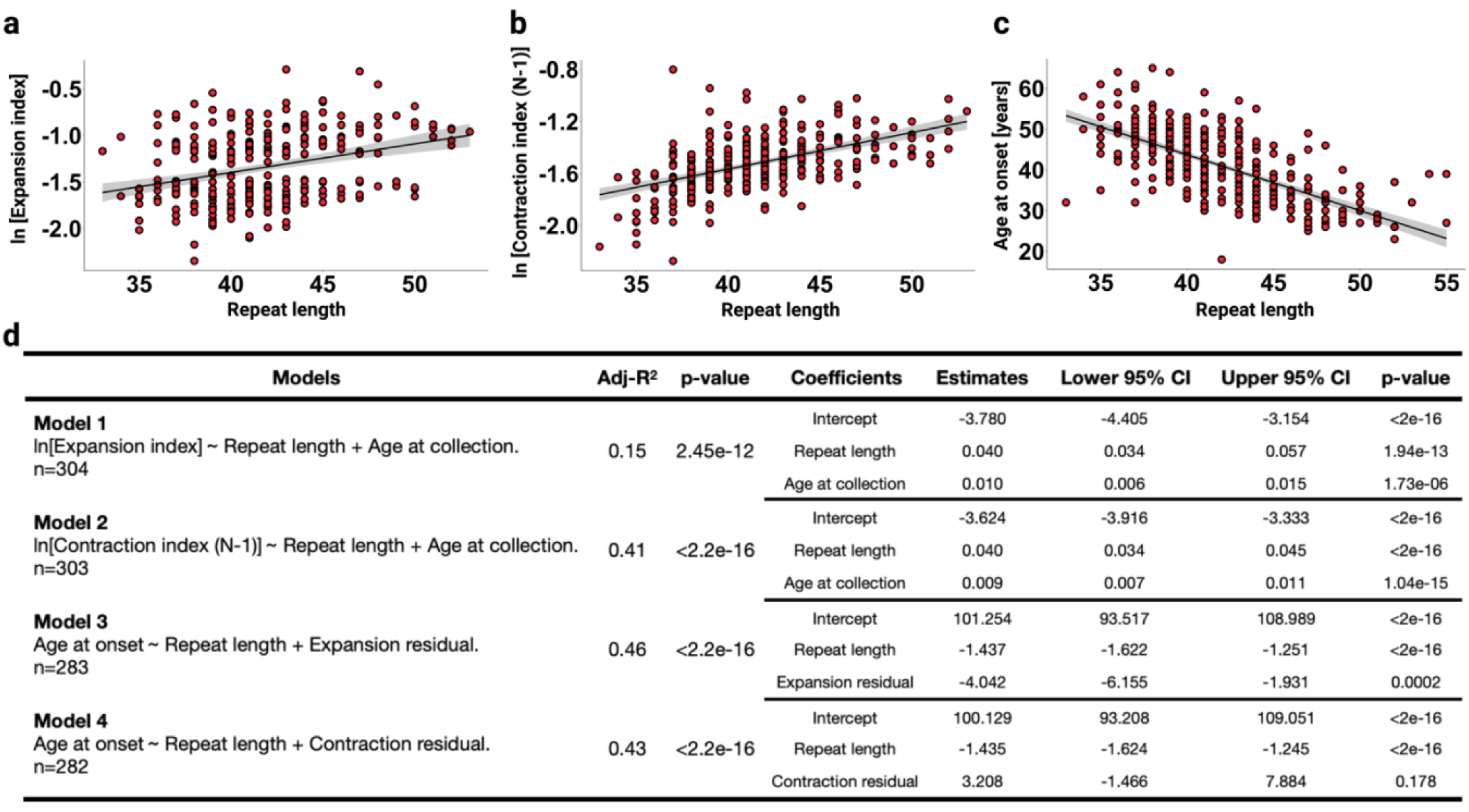
Models of CCCTCT somatic instability and age at onset in XDP blood samples. **a-c**) Correlations of somatic expansion (**a**), somatic contraction (**b**) and age at onset (AAO) (**c**) with inherited CCCTCT repeat length. **d**) Linear regression models of somatic instability (model 1 and 2) and AAO (model 3 and 4). CCCTCT instability and CCCTCT repeat length were measured by ABI fragment sizing. n=number of individuals. Each dot represents an XDP individual.

*Model 1: ln[Expansion index] ∼ Repeat length + Age at collection* (n = 304 blood samples that included age at collection)

*Model 2: ln[Contraction index (N-1] ∼ Repeat length + Age at* collection (n =303 blood samples that included age at collection)

Expansion and contraction residuals from these models were used as covariates in the following regression models of AAO:

*Model 3: AAO ∼ Repeat length + Expansion residual* (n =283 blood samples with expansion residual and AAO)

*Model 4: AAO ∼ Repeat length + Contraction residual* (n = 282 blood samples with expansion residual and AAO)

Expansion and contraction residuals were also used as phenotypes in subsequent genetic modifier analyses. For genetic analyses of AAO we derived a residual AAO accounting only for repeat length (*AAO ∼ Repeat length* (n = 377; these included 360 blood samples with AAO and 17 brain samples with AAO. Note that the modal repeat length in brain does not deviate from that in blood (ref Campion).

### Measures of somatic instability in brain tissues

Expansion and contraction indices were obtained from fragment sizing analyses, as described previously, using a 5% relative peak height threshold cut-off^28^. For genetic modifier analyses, residual expansion and contraction indices were derived in each tissue accounting for repeat length (modal allele) and age at collection*. i.e.* age at death (Expansion/Contraction index ∼ Repeat length + Age at death). Here, the full range of contraction peaks was used.

### Whole exome sequencing

DNA from 256 blood samples from XDP males was submitted for whole-exome sequencing (WES) using the Agilent SureSelect Human All Exon V6 kit (S07604514, Agilent, USA). Exome data underwent assessment of target exome coverage using Picard’s CollectHsMetrics function (https://broadinstitute.github.io/picard/). Only exomes with ≥70% of the exome covered at 10X or greater were included for downstream analyses. Whole exome data were analyzed following a multi-step quality control (QC) pipeline (**Figure S2**), with the standard Genome Analysis Toolkit (GATK) best practices recommendations^29,30^. After variant discovery with GATK, variant annotation was performed with Variant Effect Predictor (VEP V.110) using GRCh37/hg19 as a human genome reference. We subset the single nucleotide variants (SNVs) based on minor allele frequency (MAF) for single-variant and gene-based analyses using Plink V.2.0.0 (**Figure S2**). Only individuals clustering in the East Asian ancestry were considered for downstream analysis.

#### Single-variant analysis

Plink V.2.0.0 was used for the standard QC of SNVs (**Figure S2**): per-sample missing called rate (--mind) ≤ 0.1, genotype missing call rate (--geno) ≤ 0.05, Hardy-Weinberg equilibrium (HWE) = 1e-06) and MAF ≥ 0.01. We removed individuals that deviated ± 3 standard deviations (SD) from the sample heterozygosity rate mean. 248146 SNVs passed QC. We used carefully curated phenotypes (n=245 individuals for repeat expansion, n=242 individuals for repeat contraction, and n=235 individuals for AAO) for exome-wide association analyses. We estimated the association of SNVs with an XDP phenotype by fitting an exome-wide efficient mixed model using GEMMA (V.0.983)^31^.

GEMMA accounts for relatedness by generating a relatedness matrix and we used this and population stratification as covariates. Although we only analyzed exonic variants we applied a conservative genome-wide significance p-value of < 5.0e-08 for these analyses.

#### Gene-based association analysis

Plink V.2.0.0 was used for standard QC of rare SNVs (**Figure S2**): per-sample missing called rate (--mind) ≤ 0.1, genotype missing called (--geno) ≤ 0.05, and MAF < 0.01. 208707 SNVs passed QC and non-synonymous variants (missense, start loss, stop gain, and stop loss, n=47089) underwent whole-exome gene-based and candidate gene-based association analyses using burden (CMC-Wald) and SKAT-O tests, using relatedness matrix and population stratification as covariates. p-value < 3.2e-07 was considered exome-wide significant, based on the number of genes captured (n = 18128) and average number of variants in each gene (n = 8.43). In our initial test of 15 candidate genes, we incorporated a Bonferroni correction of the nominal p-value obtained, adjusting for 15 genes.

### Genome-wide association analysis

GWAS of blood expansion was performed in 163 samples for which genome-wide SNV genotyping data had been previously generated^21^. SNV QC was performed using Plink V.1.9.0: genotype missing call (--geno) ≤ 0.02, per-sample missing called rate (--mind) ≤ 0.02, deviation from mean heterozygosity ± 3 SD, and MAF ≥ 0.01. 443059 SNVs passed QC and underwent GWAS analysis using Gemma V.0.983^31^, where population stratification and a relatedness matrix were included in the linear mixed model. SNVs with a p-value < 5.0e-08 were considered genome-wide significant.

### Genotyping *MSH3* tandem repeats

DNA sequencing of the *MSH3* exon 1 polymorphic variant coding repeat region in gDNA samples from blood (n=302) or brain (n=60) was accomplished using the Illumina MiSeq Adapter Metagenomics 16S Targeted Protocol (Illumina). The MiSeq assay consists of two PCRs. PCR 1 uses forward and reverse *MSH3* exon 1 target sequences and the Illumina adaptor sequences (in bold text):

Forward: **TCGTCGGCAGCGTCAGATGTGTATAAGAGACAG**TGAGCCGATTCTTCCAGTC Reverse:**GTCTCGTGGGCTCGGAGATGTGTATAAGAGACAG**CCCAGTCCCAGACAGAAC CT

PCR 1 reaction conditions in a total reaction of 50 µl, contained 40 ng of DNA, 1X Qiagen buffer, 1X Q-Solution, 0.2 mM dNTPs. 0.5 U Qiagen Taq, 0.25 µM each forward and reverse primer. PCR cycling conditions were as follows: initial denaturation at 93℃ for 3 minutes, 33 cycles of 93℃ for 30 seconds, 60℃ for 30 seconds, 72℃ for 90 seconds, and a final extension at 72℃ for 10 minutes. PCR 1 products were cleaned up with Agentcourt AMPure XP beads with a 1X PCR products to beads ratio. PCR 2 was performed to attach the second part of the adapter and the dual barcodes using the Nextera XT Index Kit v2 Set A and D. PCR 2 reaction contained 5 µl of the cleaned-up PCR 1 product, 1X Qiagen buffer, 1X Q-Solution, 0.2 mM dNTPs, 1.25 U Qiagen Taq, 0.25 µM each forward and reverse primers (barcodes). PCR cycling conditions were as follows: initial denaturation at 93℃ for 3 minutes, 8 cycles of 93℃ for 30 seconds, 60℃ for 30 seconds, and 68℃ for 90 seconds. PCR 2 products were cleaned up with Agentcourt AMPure XP beads with a 1X PCR product to beads ratio. The final libraries were run on an Agilent Tape Station to check for concentration. Pools were made by combining 4 µl of each set of the 96 libraries and run on the Illumina Miseq using the MiSeq® Reagent Kit v3 (600 cycles) to produce 2 x 300 bp reads. Libraries were demultiplexed by Illumina MiSeq Control Software.

The repeat structure was identified by using the 15 bp flanking sequence immediately before and after the repeat sequence. If one or both flanking sequences was not found, the read was not analyzed. When both flanking sequences were found, the sequences of Read1 and the reverse complement of Read2 were read in 9 bp segments, starting at the 5’ end of the left flank and each repeat was compared to the previous repeat segment to determine the number of that specific repeat in succession. When the right flank was reached the repeat structure analysis stopped and the repeat structure was defined for Read1 and Read2 separately. Two SNVs - rs141080879(A/G) and rs1650697(G/A) - captured in the sequencing were recorded in phase with each repeat allele.

To confirm the *MSH3* repeat alleles, we also performed the fragment sizing analysis. PCR was performed using Qiagen Taq polymerase kit. The PCR reaction contained 40 ng of DNA, 1X Qiagen buffer, 1X Q-Solution, 0.2 mM dNTPs. 0.5 U Qiagen Taq, 0.25 µ M each forward and reverse primer.

PCR cycling conditions were as follows: initial denaturation at 93℃ for 3 minutes, 33 cycles of 93℃ for 30 seconds, 60℃ for 30 seconds, 72℃ for 90 seconds, and a final extension at 72℃ for 10 minutes. Primers were as follows:

Forward: 5’-[6FAM]-TGAGCCGATTCTTCCAGTC-3’ Reverse: 3’-CCCAGTCCCAGACAGAACCT-3’

The PCR products were run on the Applied Biosystems 3730xl DNA Analyzer with Genescan 500 LIZ as the internal size standard. The results were analyzed with GeneMapper v5 (Applied Biosystems).

### Haplotype analysis

We used *MSH3* repeat genotypes identified using MiSeq as above, and SNV genotypes from the exome sequencing to construct *MSH3* haplotypes. We used individuals homozygous for the three most common repeat alleles (3a/3a, 6a/6a, 7a/7a) to identify seven repeat + SNV haplotypes (3a, 6a-1, 6a-2, 6a-3, 6a-4, 7a-1, 7a-2; **Table S18**). Note that rs1650697 variant, captured in phase with *MSH3* repeats, confirmed the genotype obtained in the exome sequencing and was the major variant that distinguished haplotypes. rs141080879, also captured in phase with the *MSH3* repeat, was not informative in distinguishing haplotypes (all individuals had A allele). We included in our analyses these homozygous individuals as well as heterozyogus individuals whose genotypes could be resolved as pairwise combinations of these seven identified haplotypes.

### PacBio PCR amplicon sequencing

The CCCTCT repeat was PCR amplified using reaction conditions described above (“Determination of XDP repeat length”) from blood DNAs of 37 XDP males and an XDP patient repeat-containing bacterial artificial chromosome (BAC)^10^. PCR products from at least 5 independent PCRs were pooled and purified using the MiniElute PCR Purification Kit (Qiagen) to obtain at least 100 ng of DNA in the band containing the repeat tract (∼350 bp-550 bp), as estimated using Agilent DNA Bioanalyzer 2100. Samples were submitted to the University of Maryland Institute for Genome Science for DNA size selection using Bluepippin, followed by Single Molecule, Real-Time (SMRT) Pacific Bioscience (PacBio) sequencing. The number of highly accurate long (HIFI) Circular consensus sequence (CCS) reads obtained ranged from ∼42,000 to 93,270. Reads from patients were aligned to a custom BAC reference and displayed as waterfall plots (**Figure S3**). We developed two methods to count the number of CCCTCT (or reverse complement AGAGGG) repeats, and to accurately identify the structure of the repeat tract in each HIFI CCS read. These two analyses methods are summarized in **Figure S4**. Method 1, based on a similar pipeline to that used for analysis of *HTT* CAG MiSeq sequence^24^ (Method 1.py, See supplementary file), is unbiased to the underlying sequence and identifies all reads that begin with at least 10 CCCTCT repeats and ends with a 3’ flanking “Stop” sequence. Method 2 identifies 5’ and 3’ flanking sequences and three specific repeat structures - (CCCTCT)_n_, (CCCTCT)_n_CCT(CCCTCT)_2_ or (CCCTCT)_n_CCCT(CCCTCT)_2_ within the flanks (**Figure S4**). The script for method 2 can be found at https://github.com/alanmejiamaza. Both methods yielded similar proportions of these three sequence structures (**Table S1**).

### PacBio whole genome sequencing

30 μg blood DNA from two male XDP patients were submitted for whole genome PacBio sequencing at the University of Maryland Institute for Genome Science. Sequencing was carried out in a 30-hour movie length in 8 SMRT cells (4 cells each). Overall coverage in these two XDP samples was ∼30X (patient 1) and ∼34X (patient 2). CCS reads were assembled and mapped to the hg19/GRCh37 human genome. As the SVA is frequently observed in the human genome, only reads spanning intron 32 of the *TAF1* gene were selected for further analysis. We counted 5 (patient 1) and 9 CCS reads (patient 2) spanning *TAF1* intron 32. These CCS reads were manually evaluated for the number of CCCTCT repeats and sequence structure (Table S2).

### RNA-seq analysis

We determined the effects of *MSH3* repeat allele on *MSH3* expression levels using postmortem brain samples from 45 XDP individuals. We profiled 4 brain regions using RNASeq and performed targeted analysis of *MSH3* gene expression (caudate tissue, cerebellar cortex, BA9 frontal cortex, occipital cortex). RNA was extracted from brains after bead-based homogenization and trizol extraction (Invitrogen). Paired-end 150-bp polyA-enriched RNASeq libraries were generated (Illumina TruSeq) and sequenced on an Illumina platform (up to 50M reads per sample). After performing quality control, reads were aligned to the human genome reference (GRCh38) using STAR v2.7.10. Counts were normalized using DESeq2’s median-of-ratios method, and then technical covariates were regressed using the SVA package prior to generating expression values.

## RESULTS

### Expansion, but not contraction of the CCCTCT repeat in blood, contributes to XDP onset

We previously demonstrated repeat length-dependent and tissue-specific expansion of the XDP CCCTCT repeat that was greater in brain tissues than in blood^19^. Despite the relatively modest instability in blood DNA, it can be readily measured from fragment sizing data obtained from bulk PCR-based analyses^19^. As the number of post-mortem XDP brains is limited, here we performed our genetic discovery analyses using blood DNA from affected males to maximize sample size. Using small pool-PCR (SP-PCR) in brain tissues we also showed previously the occurrence of repeat contractions as well as repeat expansions^19^. Notably, SP-PCR sizing of single molecules revealed a single peak with marginal or absent PCR slippage products (**Figure S5**), confirming that the distribution of both contraction and expansion peaks observed in bulk PCR represents somatic contraction and expansion events, and are not artifacts of the PCR. We therefore used bulk PCR analyses of blood DNA to derive quantitative metrics of both expansion and contraction (See Materials and Methods)^19,28^.

We first set out to test whether CCCTCT expansion and/or contraction, measured in blood DNA, contribute to XDP AAO in affected males. Expansion and contraction indices were dependent on inherited repeat length (assumed to be the modal repeat detected in blood) and age at collection (**Figure 1a, b, d,** regression models 1 and 2, **Figure S1**). This larger dataset confirms our previous observation of repeat length-dependent expansion in blood^19^, extends this relationship to repeat contraction, and shows that repeat expansion and contraction accumulate with age in blood (**Figure S1)**. Importantly, the age-dependence of both expansions and contractions reinforce data from single molecule SP-PCR (**Figure S5**) that both expansion and contraction peaks in PCR products from bulk DNA represent somatic mutation events and are not simply PCR artifacts. From regression models 1 and 2 (**Figure 1d**), we derived expansion and contraction residuals (Materials and Methods), representing scores of individual-specific measures of blood repeat expansion or repeat contraction independent of inherited repeat length and age at blood collection. Then we tested the contribution of expansion residual or contraction residual to AAO in males from whom we had both blood instability and AAO data (**Figure 1d**, regression models 3 and 4). We found that both the inherited repeat length and expansion residual significantly contributed to an earlier AAO (**Figure 1d**, regression model 3), together explaining 46% of the variation of XDP onset. In contrast, contraction residual in blood did not contribute significantly to AAO (**Figure 1d**, regression model 4). Thus, these data support the contribution of repeat expansion, which can be captured in blood DNA, to XDP AAO, and suggest the existence of shared genetic modifiers of AAO and repeat expansion in XDP. The data do not rule out a contribution of repeat contraction to AAO but suggest that any potential contribution is less well captured by a readout in blood.

### Analyses of sequence variation in the CCCTCT repeat

Noncanonical repeat-containing sequences such as those containing repeat interruptions have been shown to modify disease phenotypes, *e.g.* in HD and in myotonic dystrophy type I (DM1)^23,26,32^, and a recent study reported the presence of mosaic divergent repeat interruptions of the XDP repeat tract^33^. We therefore first examined the presence of variant repeat structures in our XDP cohort by performing PacBio long-read sequencing of repeat-containing PCR amplicons from the blood of 37 males. These included 17 individuals and 14 individuals that fell into the 10^th^ percentile and 90^th^ percentile “extremes” of AAO and/or repeat expansion, respectively. We also performed PacBio amplicon sequencing of a previously sequenced XDP patient BAC^10^, as well as PCR-free whole genome PacBio sequencing of blood DNA from two XDP males. Waterfall plots showing the repeats and allele structure are shown in **Figure S3**. Using two independent alignment-free methods (Materials and Methods, **Figure S4)** we extracted sequences with either (CCCTCT)_n_, (CCCTCT)_n_CCT(CCCTCT)_2_ or (CCCTCT)_n_CCCT(CCCTCT)_2,_ structures, previously described^33^. We found that the vast majority (mean ∼ 98 % by each method) of these reads from the amplicon sequencing of all 37 blood samples contained the sequence structure (CCCTCT)_n_CCT(CCCTCT)_2_ (**Table S1**). The same structure was found in ∼97% of the reads from the BAC (**Table S1, Figure S3**), confirming the that previously determined by deep-sequencing assembly of BAC clones of the region spanning the SVA^10^. The modal repeat length [‘n” within the (CCCTCT)_n_CCT(CCCTCT)_2_ sequence] was highly correlated with that obtained by fragment sizing (**Figure S6)**. We also detected reads at low frequencies, both in the patient samples and in the BAC, with either (CCCTCT)_n_ or (CCCTCT)_n_CCCT(CCCTCT)_2_ structures (**Table S1**). PCR-free whole genome sequencing, while low depth, confirmed the predominance of (CCCTCT)_n_CCT(CCCTCT)_2_-containing reads and concordance of modal repeat length with that obtained via amplicon sequencing (**Table S2**). Therefore, in the samples examined thus far that included phenotypic extremes, we have not identified any variation between individuals in the repeat-containing structure, nor do we find evidence for within-individual variant repeat mosaicism at a level reported previously^33^. Further sequencing of our cohort would be needed to assess the presence of rarer inter-individual variation in repeat structures and to further assess intra-individual variant repeat mosaicism.

### Shared and differential impacts of *MSH3* alleles on age at onset and repeat instability in blood

Given the above observations suggesting that in our cohort, variation between individuals in the CCCTCT repeat structure may be relatively rare, we subsequently focused on the role of *trans*-acting modifiers. A previous GWAS of 353 XDP males showed significant associations of AAO with SNVs at/close to the *MSH3* and *PMS2* genes^21^. Complementing this study, and with a goal of capturing additional genetic variation that might be present in the unique XDP population of Filipino origin, we performed whole exome sequencing in a cohort 256 males (250 passing QC) (**Figure S2**). We first performed exome-wide association studies (ExWAS) with variants of ≥ 1% MAF (present in our dataset). Using residual AAO as the phenotype (AAO not accounted for by inherited repeat length: see Materials and Methods) in 235 individuals with exome and AAO data we identified a signal (signal 1) tagged by SNV rs1650697, with genome-wide significance, and which also included rs1245016 (**Table 1**, **Figure 2, Table S3, Figure S7**). rs1650697 is located within exon 1 of *MSH3* and the 5’UTR of the dihydrofolate reductase (*DHFR)* gene, with the minor (A) allele associated with earlier onset (**Table 1)**. After conditioning on rs1650697, no other variant at the *MSH3* locus remained close to genome-wide significant (**Figure S8, Table S4).** Thus, despite the relatively small cohort size we are able to identify a modifier signal at the *MSH3/DHFR* locus, recapitulating findings in the GWAS^21^. We did not detect a significant signal in *PMS2*, likely because the intronic modifier SNV detected in the GWAS^21^ is not in LD with any SNVs captured in the exome sequencing. Interestingly, rs8087221 on Chr18 emerged as genome-wide significant after the conditional analysis (**Table S4**). However, given the low minor allele frequency of this SNV (1.1%), this effect has a high likelihood of being spurious and would need further confirmation.

**Figure 2.**
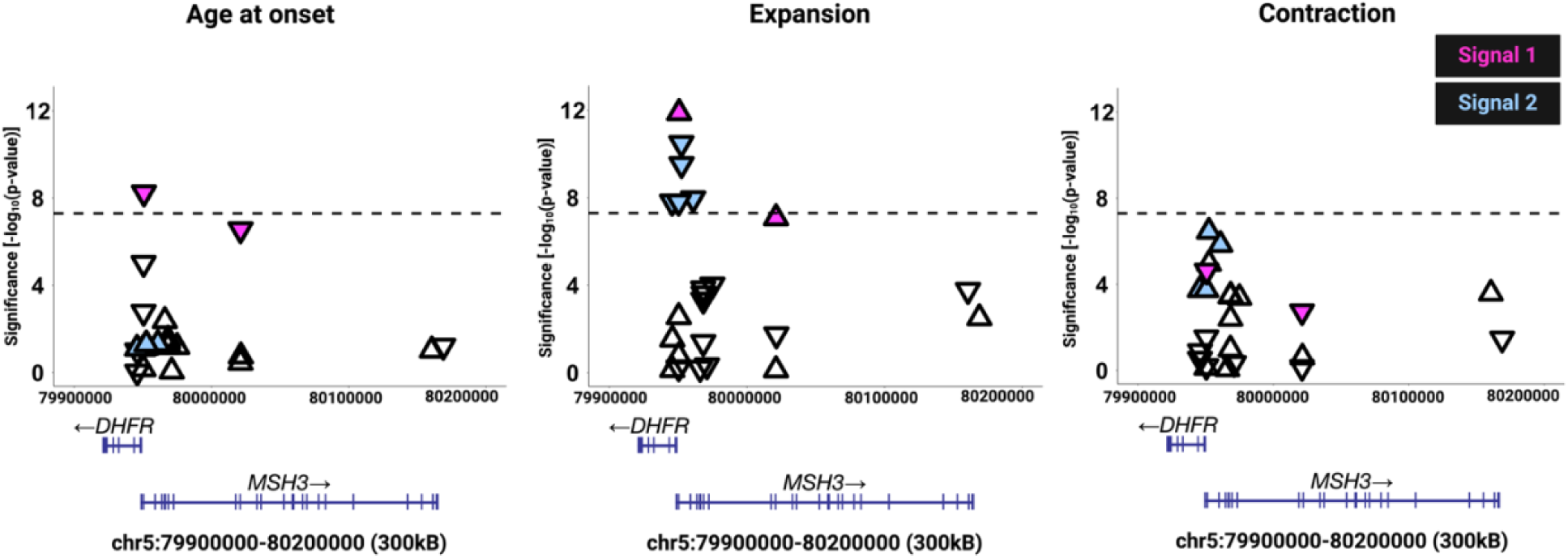
Exome-wide association analyses for age at onset and CCCTCT somatic instability in blood. ExWAS for residual AAO, residual somatic CCCTCT expansion and residual somatic CCCTCT contraction in blood, performed on SNVs with minor allele frequency ≥ 0.01, showing the region of the *MSH3/DHFR* locus in which significant associations were detected. Genomic coordinates are based on GRCh37/hg19. Signal 1 and signal 2 SNVs are indicated in pink and blue respectively and are identified as separate signals based on conditional analyses (**Figure S8, Tables S4, S6, S9**). Downward-pointing triangles show SNVs associated with a lower residual value (earlier AAO, less expansion, less contraction) and upward-pointing triangles show SNVs associated with a higher residual value (later AAO, more expansion, more contraction). The horizontal dotted lines indicate a genome-wide significance threshold of p-value = 5e-08.

**Table 1.** Top Chr 5 modifier signals in exome-wide association tests.

| rs ID | Chr:location<br>(hg19) | Chr5<br>signal | Effect<br>Allele | AF | Nominal test |  | Conditional test |  |
| --- | --- | --- | --- | --- | --- | --- | --- | --- |
|  |  |  |  |  | Beta ± SE | p-value | Beta ± SE | p-value |
| AAO |  |  |  |  |  |  |  |  |
| rs1650697 | 5:79950781 | 1 | A | 0.372 | -0.33 ± 0.54 | 5.68e-09 | - | - |
| Expansion |  |  |  |  |  |  |  |  |
| rs1650697 | 5:79950781 | 1 | A | 0.359 | 0.22 ± 0.03 | 1.31e-12 | - | - |
| rs1643639 | 5:79952390 | 2 | C | 0.112 | -0.32 ± 0.04 | 3.27e-11 | -0.24 ± 0.04 | 4.05e-08 |
| Contraction |  |  |  |  |  |  |  |  |
| rs1643639 | 5:79952390 | 2 | C | 0.114 | 1.06 ± 0.02 | 3.48e-07 | - | - |

The contribution of blood somatic expansion to AAO (**Figure 1d**, regression model 3) predicted that modifiers of AAO also modify expansion in blood. We therefore performed ExWAS using residuals of repeat expansion (derived from regression model 1, **Figure 1d**) as the phenotype in 245 individuals with exome and expansion data. We identified a genome-wide significant signal (signal 1) with rs1650697 as the top SNV, (**Figure 2**, **Table 1, Table S5, Figure S7**) and encompassing rs1245016 as in the AAO ExWAS. The minor (A) allele of rs1650697 was associated with increased somatic expansion, mirroring the direction of effect on onset and consistent with higher repeat expansion rates hastening XDP onset. Conditioning on rs1650697 revealed a distinct second genome-wide significant signal (signal 2), with the top SNV, rs1643639, within intron 2 of *MSH3* and intron 1 of *DHFR* (**Table 1, Figure S8, Table S6**). In contrast to signal 1, signal 2 minor alleles were associated with decreased expansion in blood (**Table 1**). Consistent with two distinct modifier signals, top SNVs from signal 1 and signal 2 were not in linkage disequilibrium (LD) in our XDP cohort (R^2^=0.07) nor in individuals with East Asian (EAS) background in the 1000 Genomes Project (1KG). Conditioning on both rs1650697 and rs1643639 showed no other genome-wide significant signal. We confirmed these two independent signals in a GWAS of residual expansion performed on a subset of 163 XDP males that overlapped with the previous GWAS^21^ and our exome dataset and for which genome-wide SNV genotyping data were available (**Table S7**).

To gain further insight into the somatic instability in XDP blood, we performed ExWAS on the repeat contraction residual phenotype (derived from regression model 2, **Figure 1d**) in 242 individuals with exome and contraction data. rs1643639 (top SNV for signal 2 in the repeat expansion ExWAS) gave the strongest association signal and was associated with increased contraction, though the p-value of 3.48e-07 did not reach genome-wide significance (**Figure 2**, **Table 1, Table S8, Figure S7**). Conditioning on rs1643639 revealed no other genome-wide significant signal (**Figure S8, Table S9)**.

Signal 1 did not emerge as a significant modifier of repeat contraction (p-value = 2.37e-05), consistent with this signal being the major modifier of AAO (**Figure 2**) and the lack of contribution of blood contraction to AAO (**Figure 1d**, regression model 4). Linear regression modeling showed that the lead SNVs from signals 1 and 2 accounted for 29% of the residual somatic expansion, and 16% and 0.08% of the residual AAO and somatic contraction, respectively (**Table S10**). Note that although the *MSH3* variants did not meet genome-wide significance threshold in the contraction ExWAS, these regression analyses showed that rs1650697 significantly reduced contraction (p-value=0.016), while rs1643639 significantly increased contraction (p-value=0.00065) (**Table S10**). When contraction was included as a covariate in the repeat expansion model, rs1650697 and rs1643639 remained significantly associated with expansion (p-value =3.62e-9 and p-value =4.77e-8 respectively) though the p-values were reduced by a factor of ∼10 (**Table S10**). When expansion was included as a covariate in the repeat contraction model, p-values for rs1650697 and rs1643639 were also reduced, with rs1643639 still significantly associated with contraction (p-value =0.0069), while the effect of rs1650697 did not meet statistical significance but still trended in the same direction (p-value =0.089) (**Table S10**). These observations indicate that the effects of variants rs1650697 and rs1643639 on contraction are in part independent of their effects on expansion and *vice versa*. Consistent with the directions of effect of signal 1 and signal 2 variants on the XDP AAO and instability phenotypes, GTEx *cis*-eQTL data show that rs1650697 effect allele “A” is associated with higher expression of *MSH3* in blood, cortex, and basal ganglia, while in contrast rs1643639 effect allele “C” is associated with lower expression of *MSH3* (**Figure S9**).

In summary, these data provide direct support for *MSH3* modifying both repeat expansion and contraction and indicate that *MSH3* modifier alleles have different relative impacts of effect on AAO and blood instability phenotypes.

### Rare *MSH3* coding variants are associated with less somatic expansion

We next assessed the association of rare protein-coding variants (MAF<1% in East Asians and our XDP cohort) with the XDP phenotypes. As our sample size is small, we initially took a candidate gene approach, examining genes (*MSH2, MSH6, MSH3, MLH1, PMS1, PMS2, MLH3, FAN1, LIG1, POLD1, ATAD5, RRM2B, TCERG1, MED15, CCDC82*) associated with HD clinical phenotypes and/or somatic *HTT* CAG expansion in blood or mouse models^23,34^. We identified 49 non-synonymous (NS: missense, start loss, stop gain, and stop loss)^35,36^ coding variants across these genes (**Table S11)** and performed gene-based tests of association (sequence kernel association test (SKAT) optimized (SKAT-O) and the CMC-Wald burden test) with the AAO, expansion and contraction phenotypes. Of the 15 genes tested, only *MSH3* showed a significant association in both tests between NS variants and repeat expansion after multiple test correction (Bonferroni correction for 15 genes) (SKAT-O Adj p-value = 2.3e-02, CMC-Wald Adj p-value = 8.0e-03, **Table S12)**. Although there were some nominally significant associations, none of these genes showed significant associations after Bonferroni correction with contraction or AAO (**Tables S13, S14**). All individuals (n=5), unrelated, carrying the *MSH3* variants were heterozygous for one of five variants in the coding region of *MSH3*: Ala58Val (Chr5:79950719:C:T), Arg574Trp (Chr5:80040391:C:T), Cys763Phe (Chr5:80071547:G:T), Leu911Val (Chr5:80109478:T:G) and Asp1000Glu (Chr5:80150135:T:G) (**Figure 3a, Table S11**). The Ala58Val variant was located in the intrinsically disordered N-terminal domain (NTD) and the remaining four were scattered across different protein domains (**Figure 3a, b**). Only four individuals had a clinical AAO (one was a non-manifesting carrier), and no obvious clustering of these SNVs was observed with AAO (**Figure 3c**). Interestingly, all variants, except Leu911Val, were present in individuals at or below the 90^th^ percentile of expansion residuals (lowest 10% extremes of expansion) suggesting that these SNVs are associated with the dysfunction of the MSH3 protein (**Figure 3c**). Two of these variants (Ala58Val and Arg574Trp) were found in two individuals in the 10^th^ percentile of contraction residuals (highest 10% extremes of contraction) (**Figure 3c**). We also performed exome-wide SKAT-O and CMC-Wald burden tests. These did not reveal any genes meeting exome-wide significance (p-value<3.2e-07) though we note that *MSH3* is among the most significant genes for expansion (**Table S15**). Overall, while no genes meet criteria of exome-wide significance in these rare SNV analyses, prior evidence that *MSH3* is a modifier lends support to the burden of rare NS *MSH3* SNVs contributing to lower somatic expansion. Exome or whole genome-sequencing in larger cohorts will be needed to confirm these observations.

**Figure 3.**
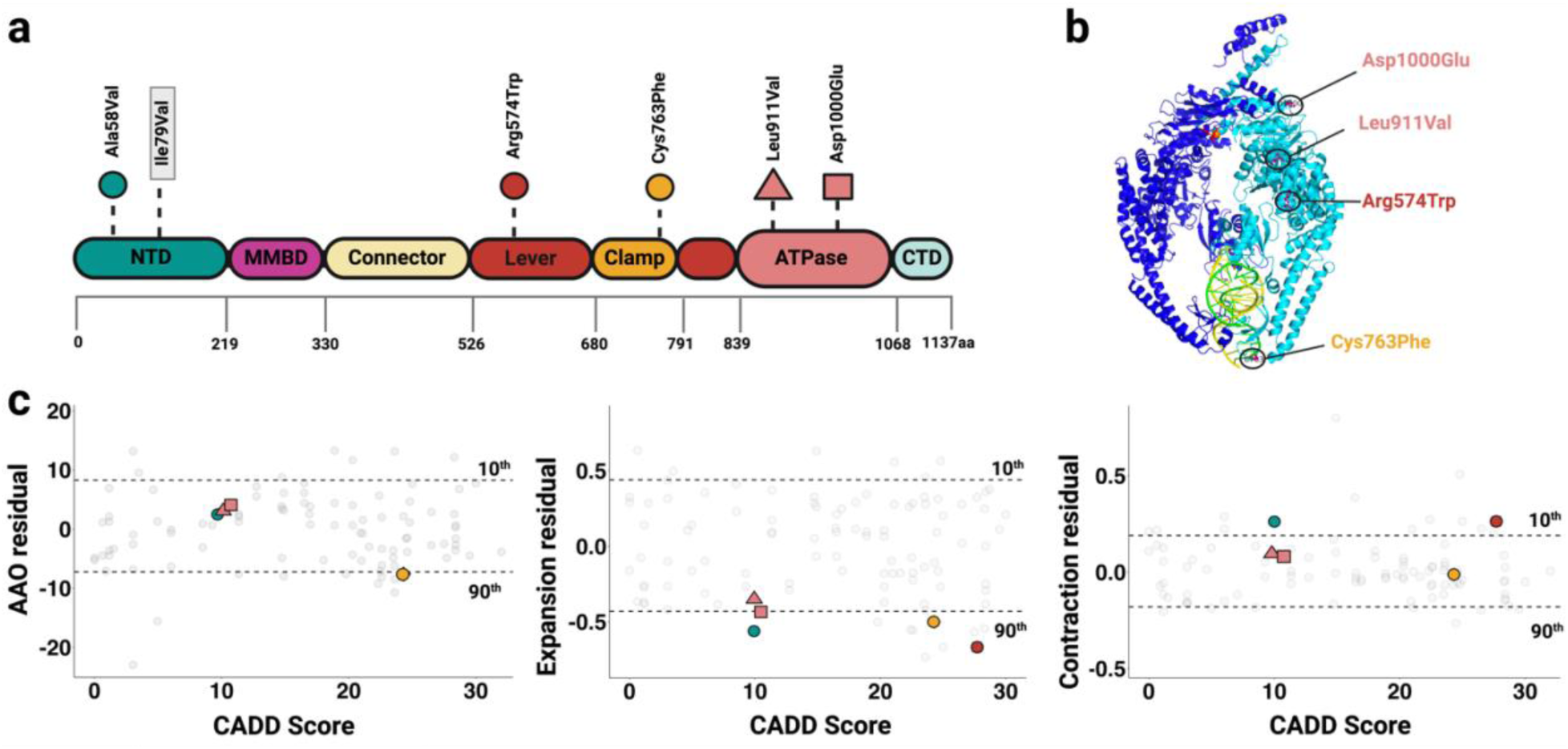
*MSH3* rare coding variants and relationships to AAO, blood expansion and contraction. **a)** Location of the five rare variants detected (Ala58Val, Arg574Trp, Cys763Phe, Leu911Val, Asp1000Glu; MAF < 0.01) and the common variant Ile79Val specified by signal 1 top SNV rs1650697 mapped onto MSH3 motifs^49^. (**b**) Three-dimensional (3D) structure of MSH3 (light blue) complexed with MSH2 (dark blue) (=MutSb dimer), modeled with PyMOL software, showing the spatial locations of the rare variants. Ala58Val, located in the unstructured N-terminal domain, is not shown. **c**) *MSH3* rare variants plotted to show their CADD score (x-axis) and AAO residual, expansion residual and contraction residuals on the y-axes. Grey circles in the background represent all data from those individuals in which non-synonymous coding changes were identified in 15 candidate genes (**Table S11**). NTD, N-terminal domain; MMBD, mismatch-binding domain, CTD, C-terminal domain. Upper and lower horizonal dotted lines show the 90^th^ and 10^th^ percentile values respectively for each phenotype. CADD scores were obtained using GRCh37-v.1.6.

### *MSH3* tandem repeat alleles modify both somatic instability and AAO

Exon 1 of the *MSH3* gene encodes a variable alanine- and proline-encoding 9 bp tandem repeat. The length of this repeat is associated with repeat expansion and clinical disease measures in HD and myotonic dystrophy type 1 (DM1) and with AAO in XDP^21,37^. To evaluate the disease-modifying effect of *MSH3* tandem repeat alleles in our XDP cohort, we performed single molecule sequencing (MiSeq) of the polymorphic repeats in 302 blood samples (**Figure 4**). The most common repeat alleles found were the 6a allele (56%), followed by the 7a (30%) and 3a (12.6%) alleles (**Figure 4a, b**). In addition, we detected novel *MSH3* tandem repeats (3c, 5c, 6d, 8c) at low frequencies (**Figure 4a, b).** Note that the 6d allele is a variant of the 6a allele that contains the Ala58Val substitution identified above. We then tested the association between the most common *MSH3* repeat genotypes (3a/3a, 3a/6a, 3a/7a, 6a/6a, 6a/7a, 7a/7a) and XDP phenotypes (**Figure 4c**, **Table S16**). We observed that 3a-containing genotypes were significantly associated with later onset, confirming previous observations^21^. We further found that 3a-containing genotypes were associated with significantly lower expansion and greater contraction (the latter not meeting statistical significance). Conversely, 6a/7a and 7a/7a genotypes were significantly associated with earlier onset, greater expansion and less contraction. Investigating the phenotypes as a function of the number of 3a, 6a or 7a alleles (**Figure S10**, **Table S17**) we found that 3a alleles significantly delayed AAO and reduced expansion, with a moderate impact on promoting contractions, while 7a alleles significantly accelerated AAO, increased expansion and suppressed contraction. The presence of two 3a alleles or two 7a alleles had a stronger modifying effect than the presence of a single allele. The number of 6a alleles had a mild impact delaying AAO and suppressing expansion.

**Figure 4.**
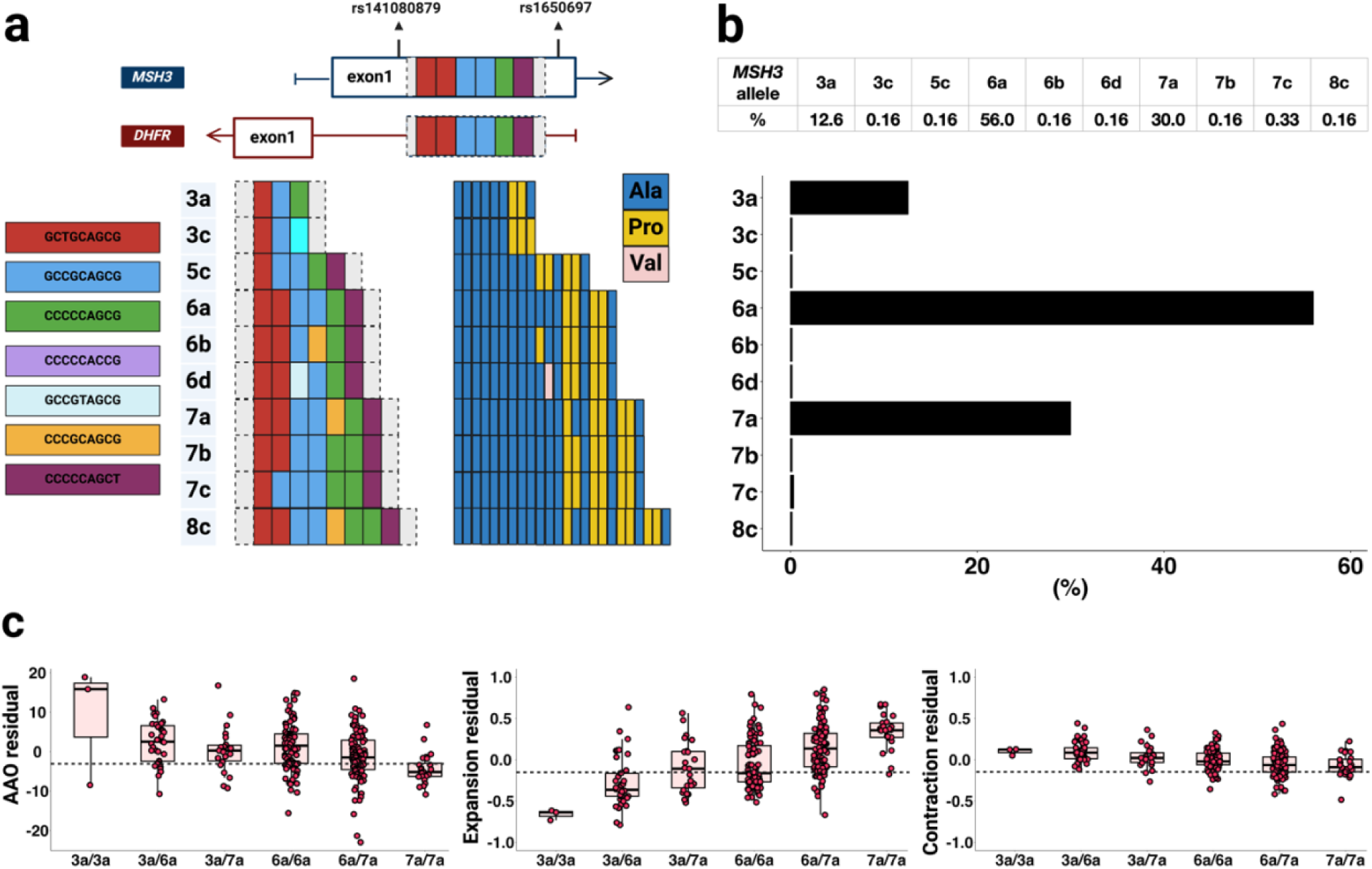
The association of *MSH3* tandem repeat variants with age at onset, blood expansion and contraction. **a**) Schematic of *MSH3* tandem repeats detected using MiSeq analyses in 302 XDP blood samples. The repeats are upstream of the *DHFR* coding region; *DHFR* shares a promoter with *MSH3* and is transcribed in the opposite direction. Each repeat variant is color-coded and the corresponding amino acids are represented. Nomenclature is according to Flower et al^37^. 6d is a variant of 6a resulting from an Ala58Val substitution. **b**) Frequency of *MSH3* tandem repeats detected. **c**) Residuals of AAO, CCCTCT expansion and CCCTCT contraction for individuals harboring 3a/3a, 3a/6a, 3a/7a, 6a/6a, 6a/7a and 7a/7a *MSH3* repeat genotypes. See **Table S16** for statistical analyses. Each dot represents an XDP individual.

We identified seven repeat haplotypes by converging SNPs from exome analysis and *MSH3* repeat alleles (Materials and Methods) and tested their effect on XDP phenotypes (**Table S18**). The only 3a haplotype identified (3a-1) included the top signal 2 SNV rs1643639 effect allele (C) and did not allow the effects (earlier AAO, less expansion, more contraction) of the 3a repeat to be distinguished from those of the rs1643639 modifier variant. Similarly, both 7a-1 and 7a-2 haplotypes included the top signal 1 SNV rs1650697 effect allele (A), tracking with later AAO, more expansion and less contraction, precluding the ability to distinguish effects due to the repeat or the SNV. Interestingly, the two most frequent 6a haplotypes, 6a-1 and 6a-2 contained the rs1650697 effect allele A and major allele G, respectively. While 6a-1 was associated with significantly earlier AAO, 6a-2 was associated with significantly later AAO (**Table S18**). Expansion and contraction phenotypes did not distinguish the effects of 6a-1 and 6a-2. These data therefore indicate, at least for AAO, an effect of rs1650697 A variant that is independent of the *MSH3* variant repeat.

### *MSH3* modifies somatic instability in brain tissues

Taken together, the results presented above support MSH3’s role in determining the timing of onset of XDP by modifying the instability of the XDP CCCTCT repeat and predict that *MSH3* modifies CCCTCT repeat instability in the brain. To test this, we derived quantitative measures of expansion and contraction in different brain tissues. We previously showed, by quantifying expansion indices from fragment sizing data of CCCTCT-containing PCR amplicons that the extent of CCCTCT expansion in brain is region- and repeat length-dependent^19^. With the knowledge from SP-PCR that contractions occur in brain^19^, we first expanded on these analyses by also quantifying contraction indices from bulk PCR fragment sizing data in a wide range of brain tissues. Interestingly, except for cerebellum, which exhibited less CCCTCT contraction than any other brain tissue analyzed, all other brain regions had relative similar contraction values (**Figure S11**). This contrasts with expansions that are more variable between brain tissues, and greatest in cortical regions^19^. Further, cerebellum exhibited less contraction than blood (**Figure S11**) but more expansion than blood^19^. Contractions also exhibited repeat length-dependence (**Figure S12**). Regression modeling in a subset of brain tissues (caudate, cerebellum, temporal pole, occipital cortex and BA9 cortex, and medial thalamus) for which we had the largest expansion and contraction datasets (31-57 individuals, depending on the region) revealed significant positive associations of repeat length and/or at death (= used as age at sampling), with expansion and/or contraction in many brain regions (**Table S19**). Power was limited in brain regions with fewer samples, and generally, repeat length and age at death appeared to be better predictors of contraction than expansion. This could potentially reflect, in part, the loss of cells with highly expanded repeats.

We then derived residuals of expansion or contraction indices based on these regression models (**Table S19)**. Given the significant associations of the *MSH3* polymorphic tandem repeat with blood expansion, contraction and AAO we then evaluated the association of residual expansion or contraction values in brain tissues with *MSH3* tandem repeat genotypes (3a/3a, 3a/6a, 3a/7a, 6a/6a, 6a/7a, 7a/7a) determined using MiSeq (**Figure 5, Table S20).** Expansion and contraction residuals from all brain tissues increased and decreased, respectively, with an increasing number of *MSH3* repeats (smallest - 3a to largest -7a), paralleling results in blood. Regression analyses (excluding the single 3a/3a individual) showed that 6a/7a and 7a/7a genotypes significantly increased expansions in all brain regions tested relative to 3a/6a (**Table S20)**. 6a/7a and 7a/7a genotypes significantly decreased contractions in BA9 cortex, relative to 3a/6a **(Table S20**). Assessing the impact of individual repeat alleles (**Table S21**), the 3a repeat had the strongest and statistically significant impact on decreasing expansion in the caudate; 3a alleles accounted for 44.6% of the residual expansion (**Table S21**). The 7a allele had the strongest and statistically significant impact on increasing expansion in the cerebellum; 7a alleles accounted for 24.1% of the residual expansion (**Table S21)**. In a subset of XDP individuals and brain regions (caudate, cerebellum, occipital cortex and BA9 cortex) we also analyzed MSH3 expression levels obtained from RNA-seq data. We observed that individuals with 3a/6a and 6a/6a genotypes tended to have the with lowest levels of MSH3 mRNA in these brain regions, while with 6a/7a or 7a/7a genotypes had higher MSH3 mRNA levels respectively (**Figure 6, Table S22**). The effect of the *MSH3* repeat alleles on MSH3 expression was particularly pronounced in caudate, where one or two 7a alleles significantly increased expression 1.1-fold or 1.3-fold, respectively (**Table S23**). These data are consistent with the association of the 3a allele with reduced MSH3 expression in blood^37^, and demonstrate correlations between MSH3 expression, more expansion and less contraction in XDP brains. Overall, we provide evidence that *MSH3* modifies the onset of XDP by altering the instability of the CCCTCT repeat in the brain.

**Figure 5.**
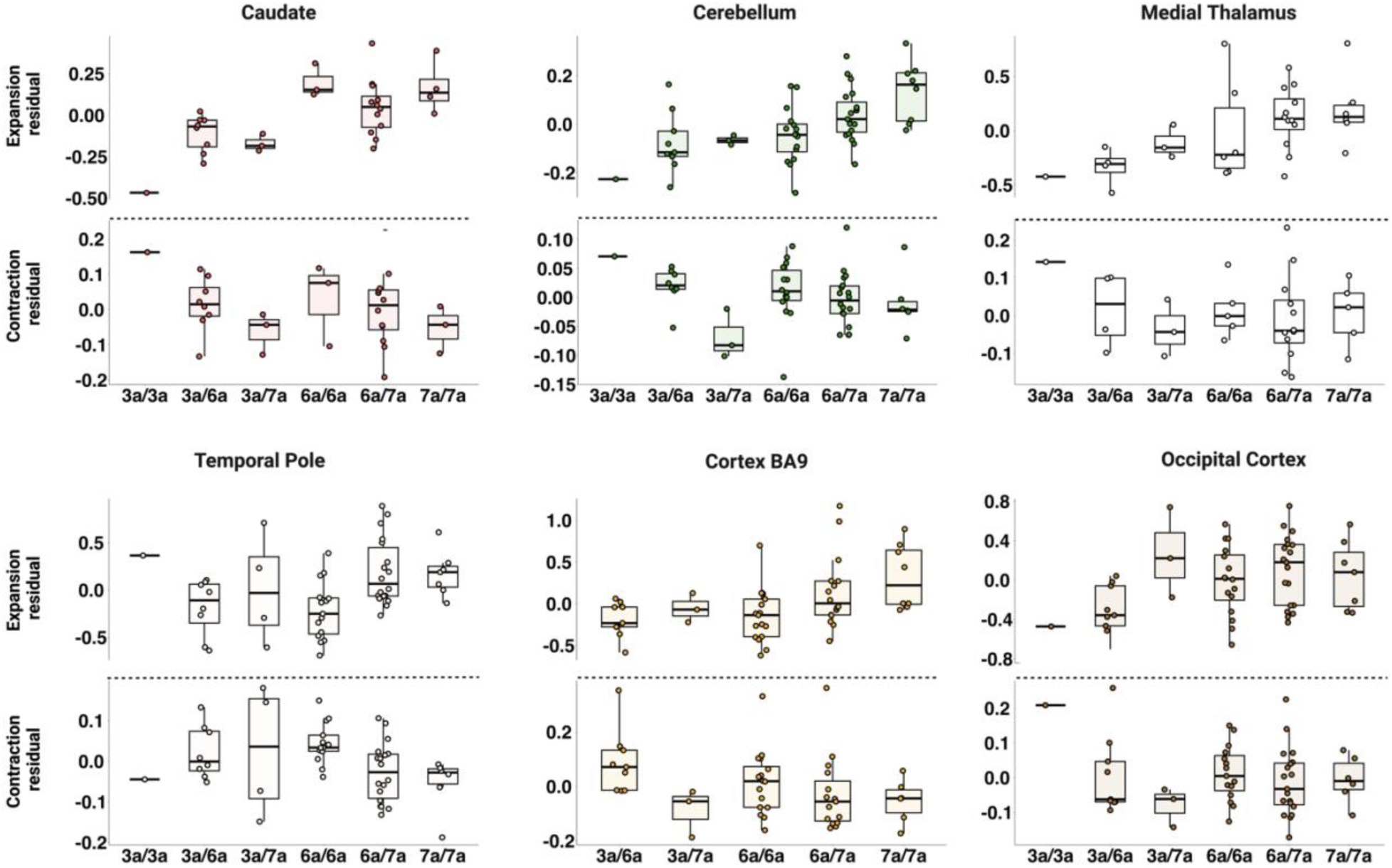
*MSH3* tandem repeat variants in XDP post-mortem brain tissues. Residuals of CCCTCT expansion (above dotted lines) and CCCTCT contraction (below dotted lines) in caudate, cerebellum, medial thalamus, temporal pole, cortex BA9 and occipital cortex for individuals harboring 3a/3a, 3a/6a, 3a/7a, 6a/6a, 6a/7a and 7a/7a *MSH3* repeat genotypes. See **Table S20** for statistical analyses. Each dot represents an XDP individual.

**Figure 6.**
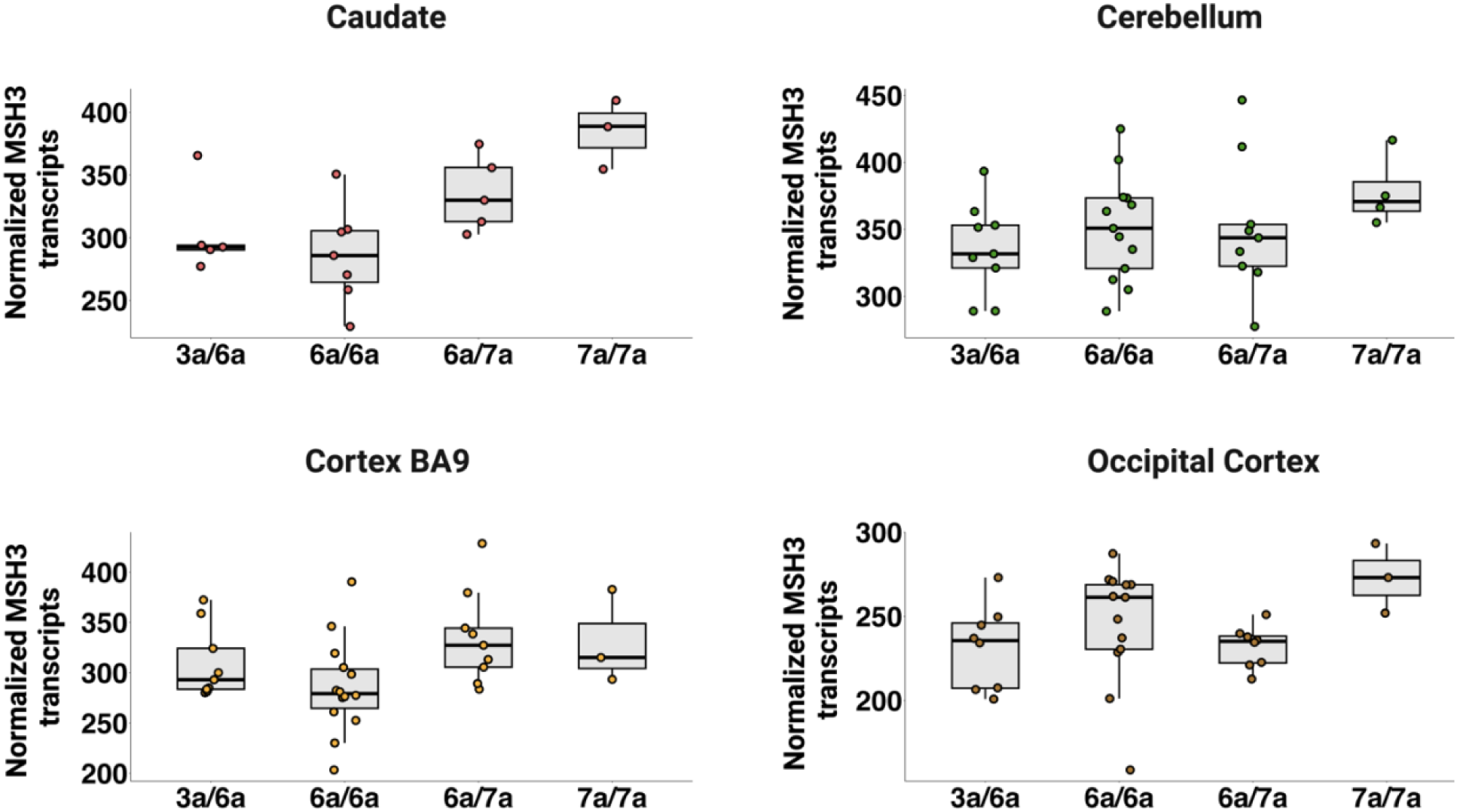
*MSH3* tandem repeat variants alter MSH3 expression levels in the brain. MSH3 mRNA expression levels in caudate, cerebellum, cortex BA9 and occipital cortex for individuals harboring 3a/6a, 6a/6a, 6a/7a and 7a/7a *MSH3* repeat genotypes. See **Table S22** for statistical analyses. Each dot represents an XDP individual.

## Discussion

Previous observations that 1) the length of the XDP SVA CCCTCT repeat tract is inversely correlated with AAO, 2) the CCCTCT repeat exhibits length-dependent somatic expansion in the brain and 3) variants in MMR genes modify AAO, support a model in which somatic CCCTCT repeat expansion drives the rate of XDP onset. Here, we demonstrate that *MSH3* variants modify somatic CCCTCT repeat instability, both in blood and in the brain. We provide the first empirical evidence for an association of *MSH3* with repeat instability in the brain in any repeat expansion disease and our findings provide key evidence that the role(s) of MSH3 in CCCTCT repeat dynamics underlie its impact on clinical disease.

We previously showed that while somatic instability of the XDP CCCTCT repeat was expansion-biased, both of the instability phenotypes (expansion and contraction) could be detected. With this knowledge, and the observation from analysis of single molecules that this hexanucleotide repeat exhibits little to no (typically contraction-biased) confounding PCR slippage artifact, we quantified both repeat expansions and contractions for integration with genetic data. In blood and brain tissues, both expansions and contractions exhibited dependence on repeat length and on age at sampling (blood) or age at death (brain), indicating somatic events that increase over time as XDP individuals age. In the brain, contractions were distinguished by an apparent lack of brain-regional specific differences that is observed for expansions, with the exception of the cerebellum that exhibits low levels of both expansion and contraction relative to other brain regions. These findings suggest that the factors (as yet unknown) contributing to cell type-specific expansion propensities and hence the levels of expansion that are reflected in tissues do not influence contractions in the same way. It is also possible that there are in fact subtle differences in contractions between brain regions that cannot be readily resolved from the bulk PCR amplicon data, as suggested by our previous single molecule analyses^19^.

In the ExWAS analysis we detected two distinguishable major modification signals, captured to different degrees by AAO and repeat expansion or contraction in blood. Signal 1, tagged by rs1650697, was associated with an earlier AAO and increased expansion in blood. Signal 2, tagged by rs1643639, was not associated with AAO, but was associated with decreased expansion and increased contraction in blood. Although the impact on contraction did not meet the criterion of genome-wide significance (potentially conservative in the context of only the exome variants analyzed in this study), the p-value of 3.48e10^-7^ was highly suggestive of a real effect. The different relative impacts of these two variants on AAO, blood expansion and contraction are unlikely to be due to differences in the power to detect modifier effects given the similar sample sizes for each analysis (AAO n=235, expansion n=245, contraction n=242). Rather, these data suggest that there may be cell type-dependent modifier effects of these *MSH3* alleles. As examples (refer **to Figure 2**) – *(i)* Relevant to AAO, signal 1 might be predicted to exert a relatively strong impact on repeat expansion compared to signal 2 in the cell type(s) in the brain that are relevant to disease onset; *(ii)* Relevant to blood contraction, signal 2 may have a relatively strong impact compared to signal 1 in the hematopoetic cells driving repeat contraction observed in blood. As both signal 1 and signal 2-tagging SNVs are eQTLs influencing *MSH3* expression levels, these differences may reflect cell type-specific eQTL effects, as suggested based on similar observations in HD GWAS^23^. To ascertain the relationship between our modifier signals and those in previous GWAS we determined the linkage disequilibrium (LD) between the top signal 1 and signal 2 SNVs with those *MSH3* SNVs identified in GWAS as modifiers of XDP AAO^21^ and those that modified either blood expansion or clinical phenotypes in HD^23^ (**Table S23**). This indicates that our signal 1 partially captures the Laabs et al. top *MSH3* SNV (rs245013) (LD between signal 1 top SNV rs1650697 and rs245013 = 0.399 in Europeans and 0.659 in East Asians). For HD, our signal 1 captures the same signal as the “5AM1” disease-hastening and “5ABEM1” blood expansion-promotion effects (LD between signal 1 top SNV rs1650697 and 5AM1 top SNV rs245100 or 5ABEM1 top SNV rs245105 = 1 in Europeans and 0.956 in East Asians). Our signal 2 partially captures the Laabs et al. rs33003 SNV that modified XDP AAO independently of rs245013 (LD between signal 2 top SNV rs1643639 and rs33003 = 0.522 in Europeans and 0.144 in East Asians). For HD, our signal 2 partially captures the same signal as the “5AM3” disease-delaying (LD between signal 2 top SNV rs1643639 and 5AM3 top SNV rs6151716 = 0.544 in Europeans, unavailable in East Asians) and “5ABEM3” blood expansion-suppressing effects (LD between signal 2 top SNV rs1643639 and 5ABEM3 top SNV rs1650689 = 0.295 in Europeans, 0.737 in East Asians). These analyses may be limited by lack of specific haplotype information in the Filipino population, nevertheless, overall, the data indicate that *MSH3* variants tagging effects on clinical phenotypes and repeat instability have shared effects across these two diseases.

Analyses of the variant alanine- and proline-coding repeat in exon 1 of *MSH3* revealed an association of the 7a repeat with an earlier AAO and with increased repeat expansion and decreased repeat contraction in both blood and in brain. In contrast, the 3a repeat was associated with a later AAO, decreased repeat expansion and increased repeat contraction in both blood and in brain. The effects of these alleles on expansion and contraction are consistent with their impact on clinical onset – *i.e*. more expansion and less contraction are associated with hastened onset, while and less expansion and more contraction are associated with delayed onset, and with the association of the variants with *MSH3* expression in brain. Haplotypes 6a-1 and 6a-2, distinguished by signal 1 SNV rs1650697, have opposite effects on onset, consistent with the onset-hastening effect of the A-allele. **Figure 7** summarizes the results of our *MSH3* common SNV and repeat variant association analyses.

**Figure 7.**
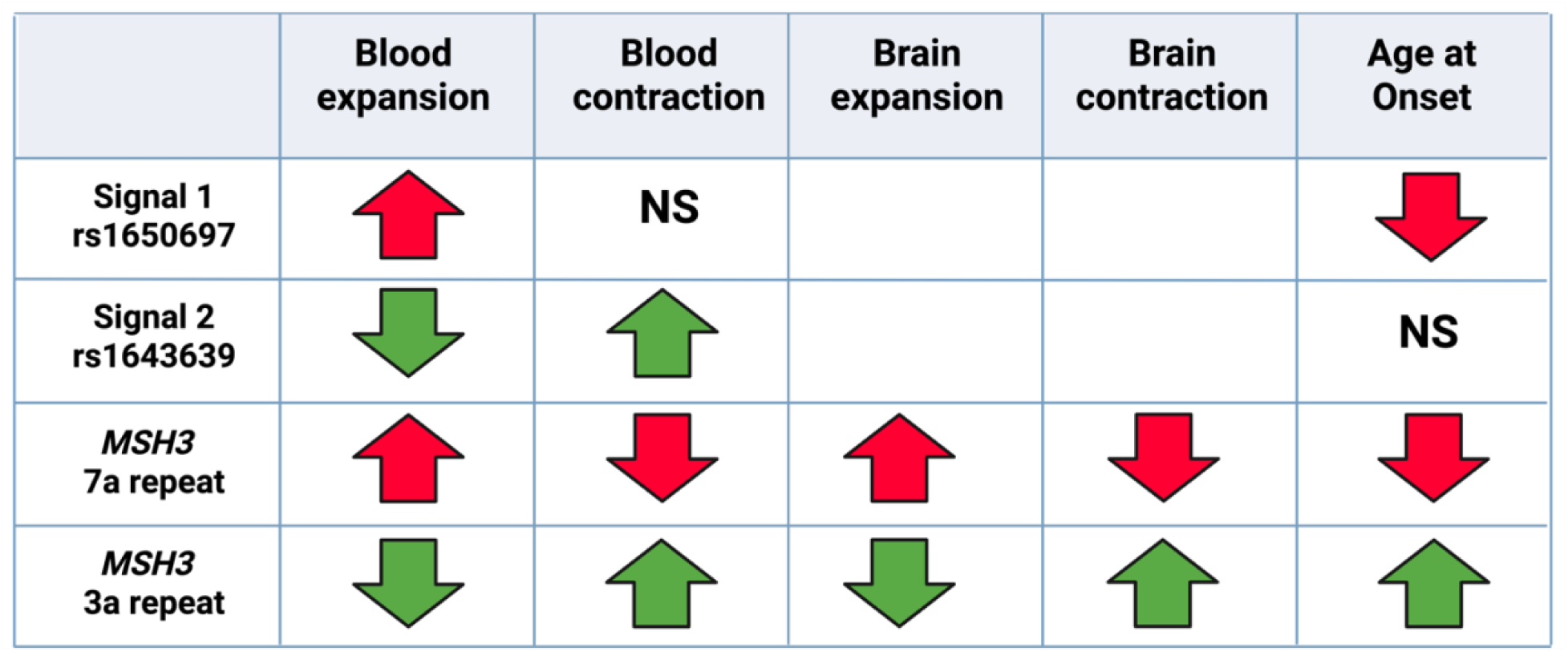
Summary of *MSH3* associations. Upward arrows indicate associations with greater residual values (more expansion, more contraction, later onset and downward arrows indicate associations with lower residual values (less expansion, less contraction, earlier onset). Red is consistent with a detrimental effect and green is consistent with a beneficial effect. SNV summaries are based on meeting genome-wide significance in ExWAS data with the exception of blood contraction signal 2 where the p-value for association was 3.48e10^-7^ (**Tables S3- S6, S8, S9**). *MSH3* repeat allele summaries are based on nominal significance in regression analyses (**Tables S16, S17, S19, S20**).

As our sample size was relatively small, we were underpowered to detect exome-wide significant effect of rare variants in gene burden-based analyses. However, *MSH3* did emerge as one of the more significant modifiers of repeat expansion in these analyses, with five non-synonymous variants found in individuals exhibiting ∼ the lowest 10% of expansion values in blood. Further understanding of the potential role of these variants will require their phasing to the common *MSH3* SNV and repeat modifier variants and studies of their impacts on MSH3 function or stability. We note that all five variants are predicted to lower MSH3’s stability (http://mupro.proteomics.ics.uci.edu/).

Taken together, our data provide strong support for somatic expansion being a major driver of XDP onset. What is the role of somatic repeat contraction in determining the onset of XDP? Although blood contraction did not predict AAO (**Figure 1**), consistent with the lack of the genome-wide significant SNVs contributing to both AAO and contraction in blood (**Figure 2**), the effect of the AAO-associated *MSH3* variant repeats on contraction both in blood and in brain (**Figures 4,5**) support an involvement of contraction. *i.e*. more contraction delays onset and *vice versa*. Based on computational modeling of repeat instability in DM1 and HD it has been suggested that expansions, which predominate in somatic cells and increase over time, are the net result of frequent expansion and contraction events with expansion rates slightly exceeding contraction rates^38,39^. Thus, the two events may be intimately coupled, and the balance between them may be influenced by different *cis-*acting (*e.g*. repeat sequence, flanking sequence) and *trans*-acting factors. In HD and in other repeat expansion diseases, expansions are thought to be driven by the binding of MutSβ (MSH2-MSH3) complex binding to repeat loop-outs^37,40,41^. The processing of such a loop-out to incorporate an expansion versus a contraction is likely to depend in part on downstream events that involve other MMR components, *e.g.* MLH3 and PMS2 that have different endonucleolytic cleavage propensities that could favor expansions over contractions^23,41,42^. Though not investigated here, further studies will be needed to understand the role of XDP onset modifier *PMS2* in CCCTCT repeat instability and to dissect the various other factors that contribute to repeat expansion and contraction of the CCCTCT repeat. Our data suggest that CCCTCT loop-outs are also targets for MSH3 binding and that higher levels of MSH3 might tip the resolution of such structures in favor of expansions over contractions, while lower MSH3 levels might favor contraction events. Interestingly, in a DM1 mouse model^43^, and to a lesser extent in an HD mouse model^44^ knockout of *Msh3* both suppressed expansions and promoted contractions in the germline. Knockout of *MSH3* also appeared to promote contractions in a human RPE1-based model of *HTT* CAG instability^45^. It is also possible that expansions and contractions might predominate in different cell types. Therefore, ultimately, resolving differential genetic modifier effects on expansions and contractions in individual cell types in the brain and in blood will be needed. Insight into the functional impacts of rare coding *MSH3* variants may also provide future opportunities to distinguish mechanisms of expansion and contraction.

Building on a framework in HD^24,34,46^ for the life-time of a vulnerable neuron. and as predicted to be applicable across many repeat expansion disorders^47^ we propose that in XDP the CCCTCT repeat must reach a threshold length(s) in target cell type(s) in order to elicit neuronal demise and ensuing clinical disease. The CCCTCT repeat can undergo somatic expansion and somatic contraction and the rate of both of these events may contribute to the timing of disease onset. Our data demonstrate a critical role for onset modifier *MSH3* in both driving expansions and protecting against contractions. Genetic variation in *MSH3* has to date also been associated with clinical disease and/or blood somatic expansion in HD, DM1and in *FMR1* premutation carriers^23,24,26,37,48^. Strategies to target MSH3 are therefore likely to have wide applicability across repeat expansion diseases. Given the genetic validation in patients and strong impact on disease onset, *MSH3* is thus a compelling therapeutic target in XDP, with strategies to lower or inhibit *MSH3* predicted to both slow CCCTCT expansion and promote CCCTCT contraction, impacting the disease course prior to clinical onset.

### Data and code availability

The datasets used and/or analyzed during the current study are available from the corresponding authors upon reasonable request. Requests for tissue specimens may be directed to. Figures and schematics were created using Biorender. Plots were done in R/RStudio V.1.3 (https://cran.r-project.org/mirrors.html).

## Acknowledgments

We thank Dr. Konrad Karczewski for his advice on exome analyses and Drs. Marc Ciosi and Darren Monckton for their inputs on modeling repeat instability. This work was supported by the CCXDP (V.C.W, L.J.O, D.C.B, M.E.T, N.S), NIH grants R01-NS102423 (D.C.B & M.E.T), R01-NS049206 (V.C.W) and R01-NS091161 (M.E.M) and the MGH executive committee on research (MGH ECOR) (A.M.M.).

## Author contributions

Conceptualization: A.M.M, V.C.W and L.J.O.; Methodology: A.M.M, A.D, and Investigation: A.M.M, M.H, K.C, T.G, A.N, A.D, R.Y, and P.D.V.M; Data analysis: A.M.M, A.D, M.H, K.C, T.G and V.C.W; Visualization: A.M.M, K.C and A.D; Resources: E.B.P, M.G.M, J.S.H, E.P.N, C.F.C, G.P.L, M.S, E.M, M.C.A, and C.C.E.D; Supervision: V.C.W and L.J.O, M.E.M, D.C.B and M.E.T; Writing – original draft: A.M.M, V.C.W and L.J.O.

**Declaration of Interests**

V.C.W. was a founding scientific advisory board members with a financial interest in Triplet Therapeutics Inc. Her financial interests were reviewed and are managed by Massachusetts General Hospital (MGH) and Mass General Brigham (MGB) in accordance with their conflict-of-interest policies. V.C.W. is a scientific advisory board member of LoQus23 Therapeutics Ltd. and has provided paid consulting services to Acadia Pharmaceuticals Inc., Alnylam Inc., Biogen Inc., Passage Bio, Rgenta Therapeutics and Ascidian Therapeutics

## Supplemental Information Supplementary figures

**Figure S1.**
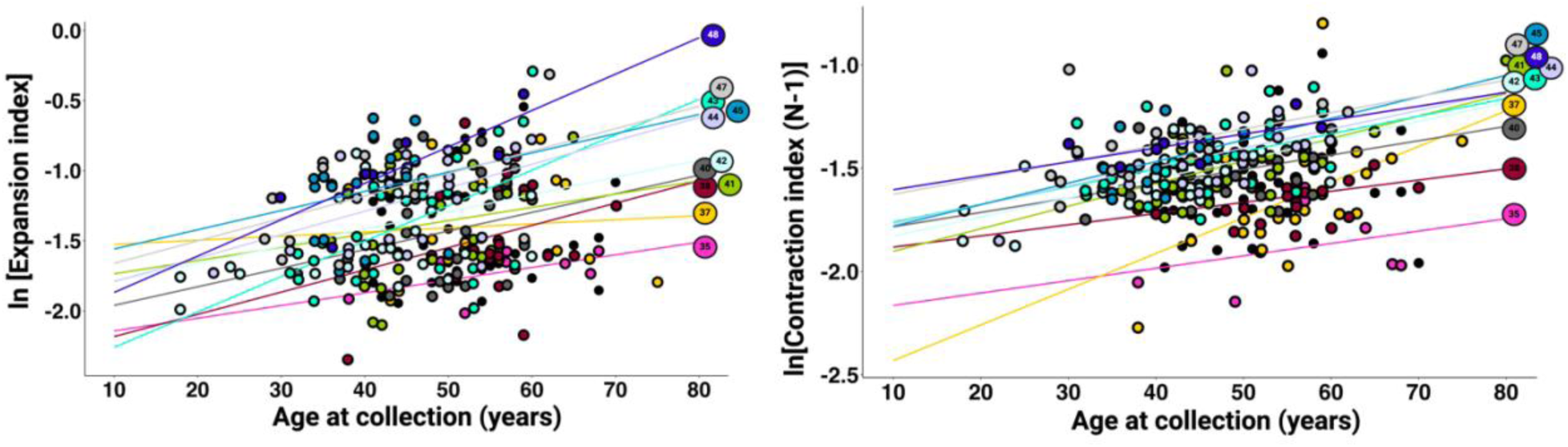
Relationship between CCCTCT expansion and contraction with age at blood collection and repeat length. Left: CCCTCT expansion, right: CCCTCT contraction. Inherited CCCTCT repeat lengths are color-coded. The graphs show that both blood expansion and contraction increase with both age and inherited repeat length and indicate a weak interaction between the two.

**Figure S2.**
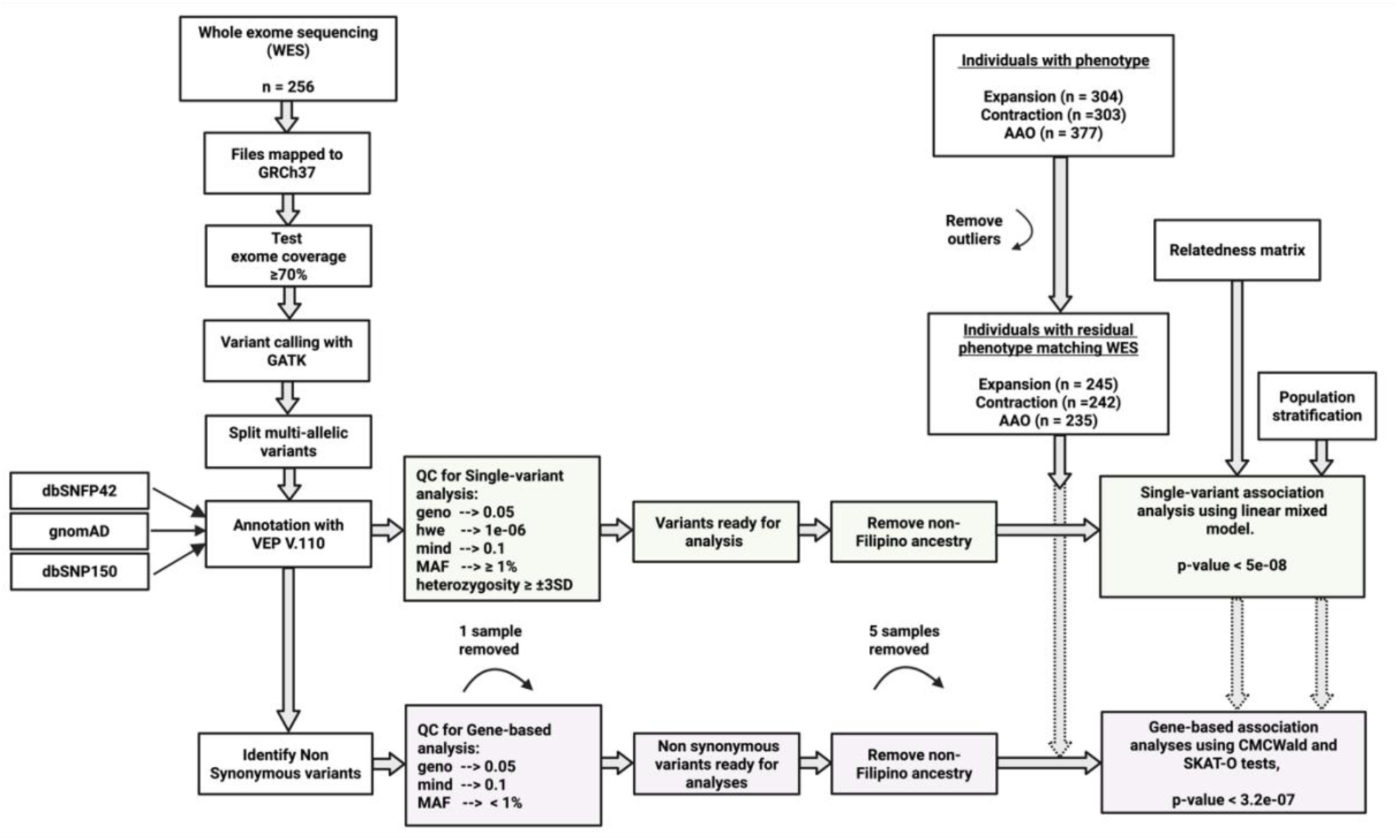
Schematic of whole exome sequencing analyses. Outline of the steps used for analyses of exome sequencing data

**Figure S3.**
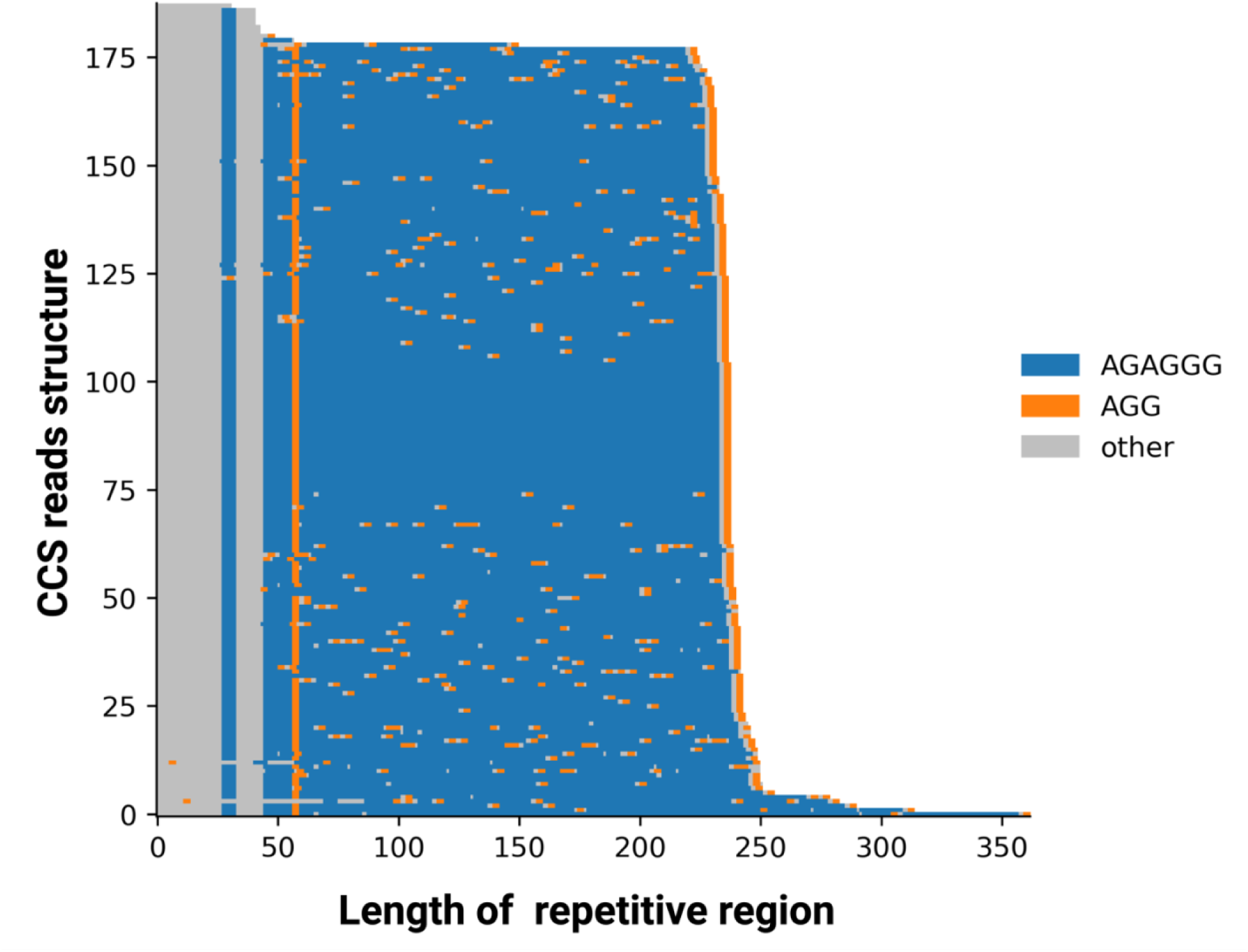
Waterfall plots of CCCTCT repeat size and structure. See Materials and Methods for details.

**Figure S4.**
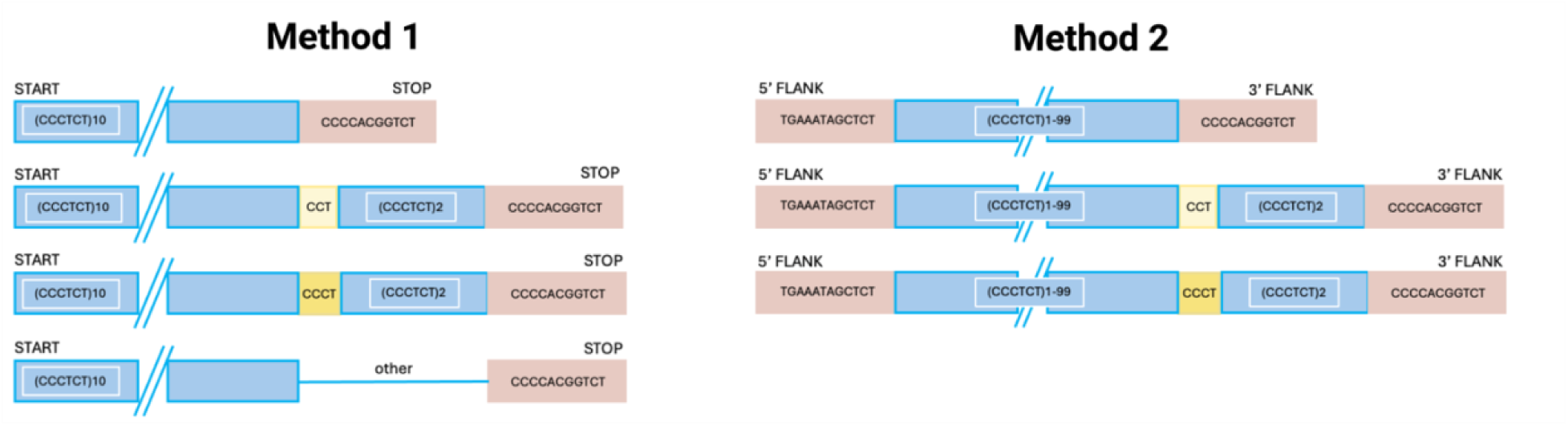
Schematic of alignment-free methods to analyze CCCTCT repeat sequences. See Materials and Methods for details.

**Figure S5.**
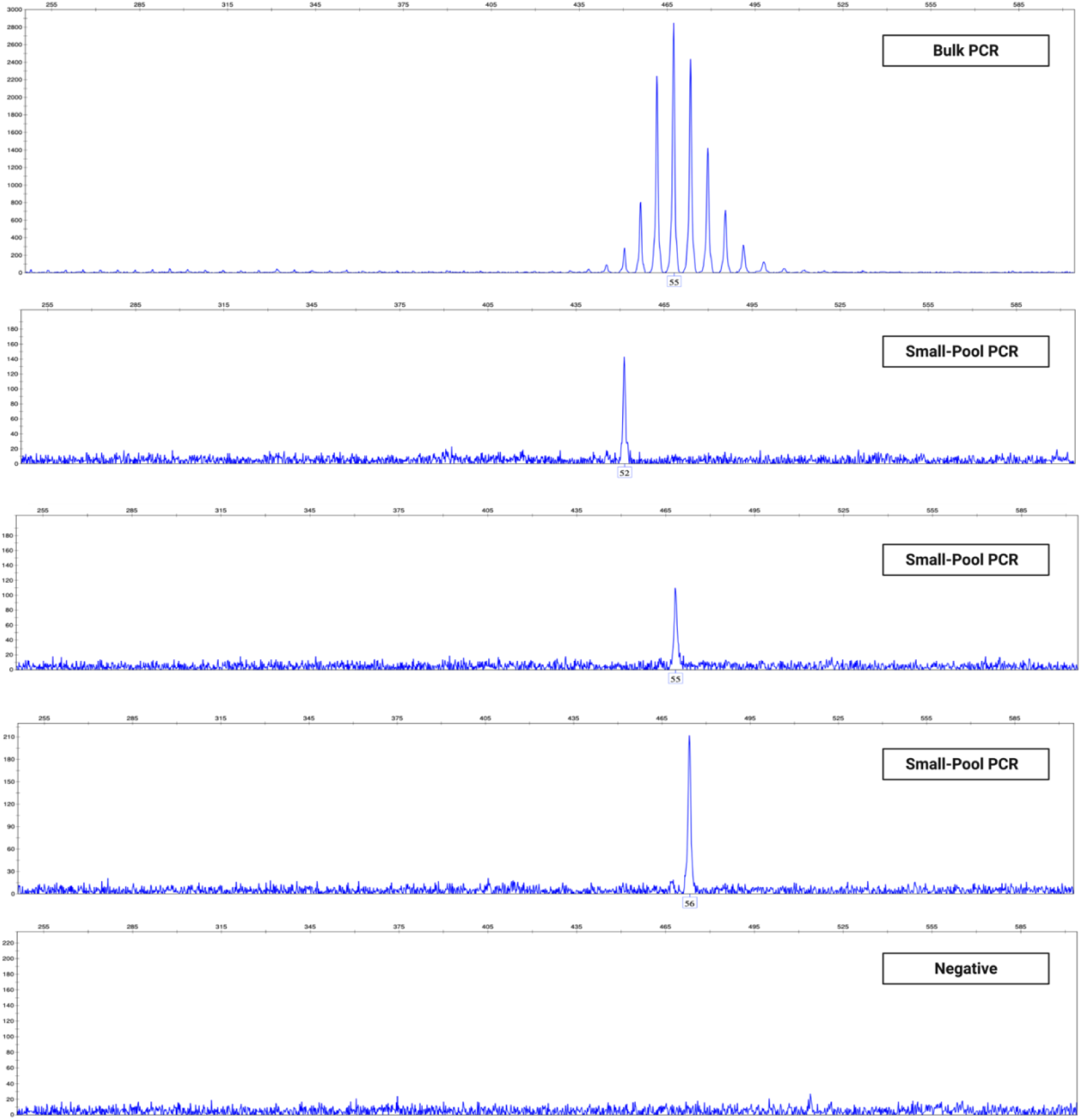
Example of small pool-PCR CCCTCT repeat amplicons. Top: Bulk fragment analysis, bottom: small-pool PCRs. The graphs display the profile for cerebellum from the same individual.

**Figure S6.**
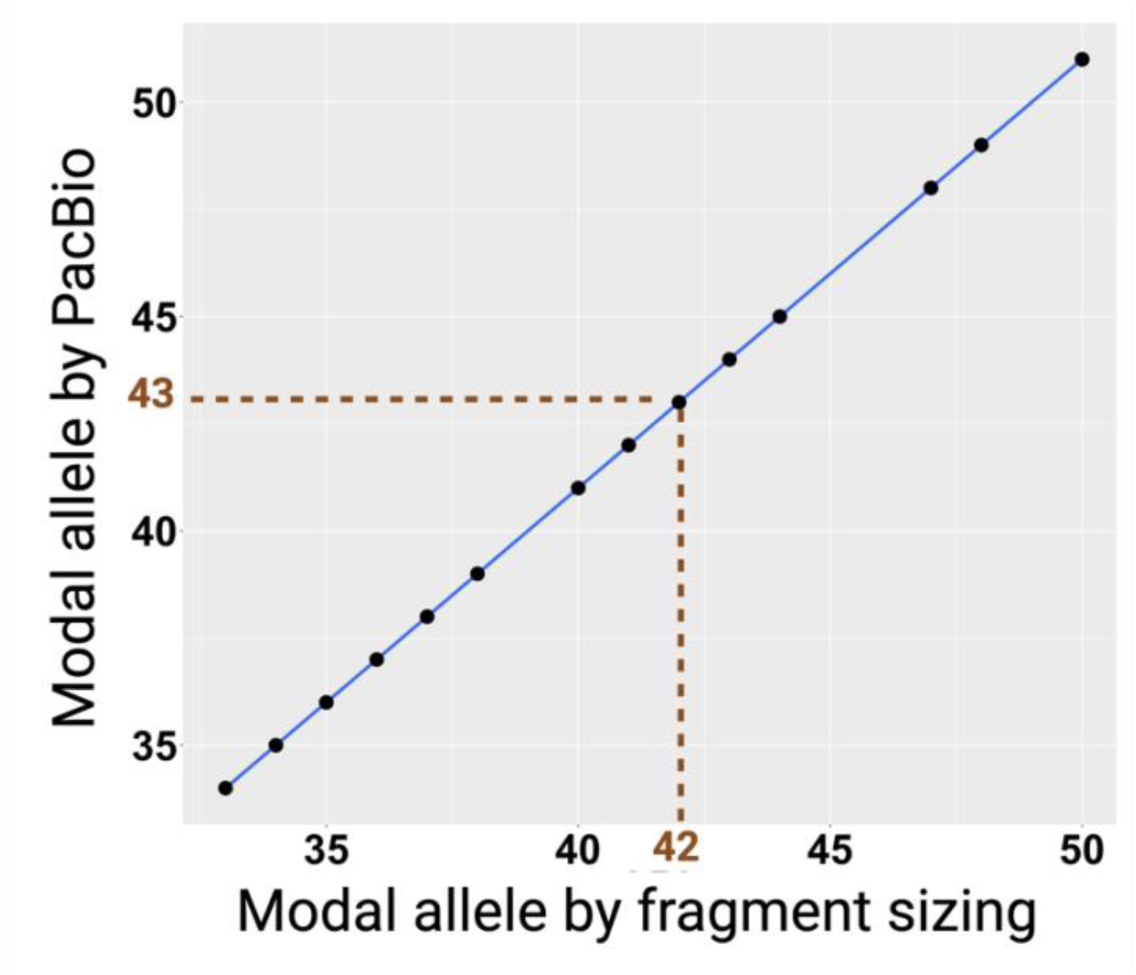
Correlation between modal CCCTTC repeat lengths obtained by PacBio amplicon sequencing and ABI-based fragment sizing. Plot of modal repeat lengths obtained by the two methods for 37 patient blood samples and a BAC clone.

**Figure S7.**
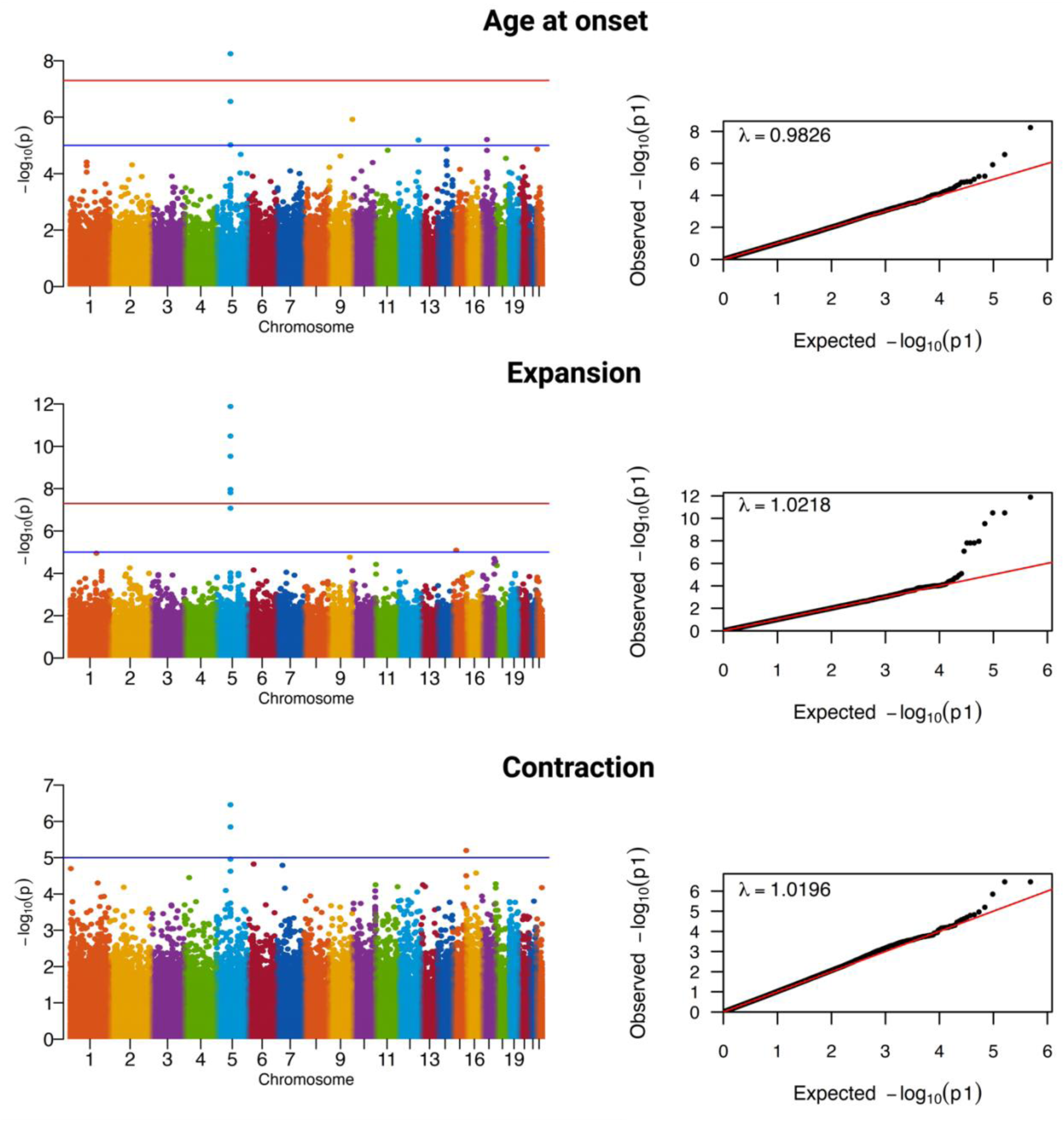
Manhattan and Q-Q plots for AAO, expansion and contraction ExWAS. See Materials and Methods for details.

**Figure S8.**
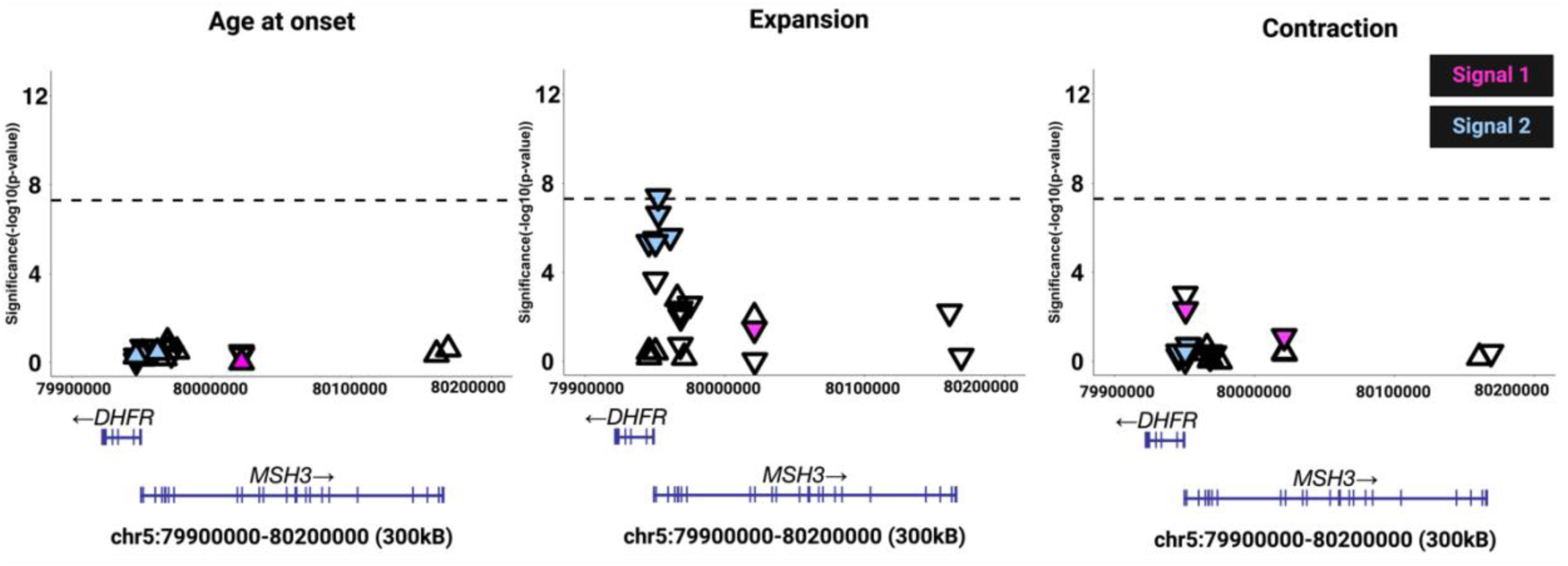
Conditional exome-wide association analyses for age at onset and CCCTCT somatic instability in blood. Conditional ExWAS for residual AAO, residual somatic CCCTCT expansion and residual somatic CCCTCT contraction in blood, performed on SNVs with minor allele frequency ≥ 0.01, showing the region of the *MSH3/DHFR* locus in which significant associations were detected. The initial ExWAS (Figure 2) were repeated, conditioning on the top SNVs for each association (rs1650697 for AAO and expansion, and rs1643639 for contraction). Genomic coordinates are based on GRCh37/hg19. Signal 1 and signal 2 SNVs are indicated in pink and blue respectively and are identified as separate signals based on these conditional analyses. Downward-pointing triangles show SNVs associated with a lower residual value (earlier AAO, less expansion, less contraction) and upward-pointing triangles show SNVs associated with a higher residual value (later AAO, more expansion, more contraction). The horizontal dotted lines indicate a genome-wide significance threshold of p-value = 5.0e-08.

**Figure S9.**
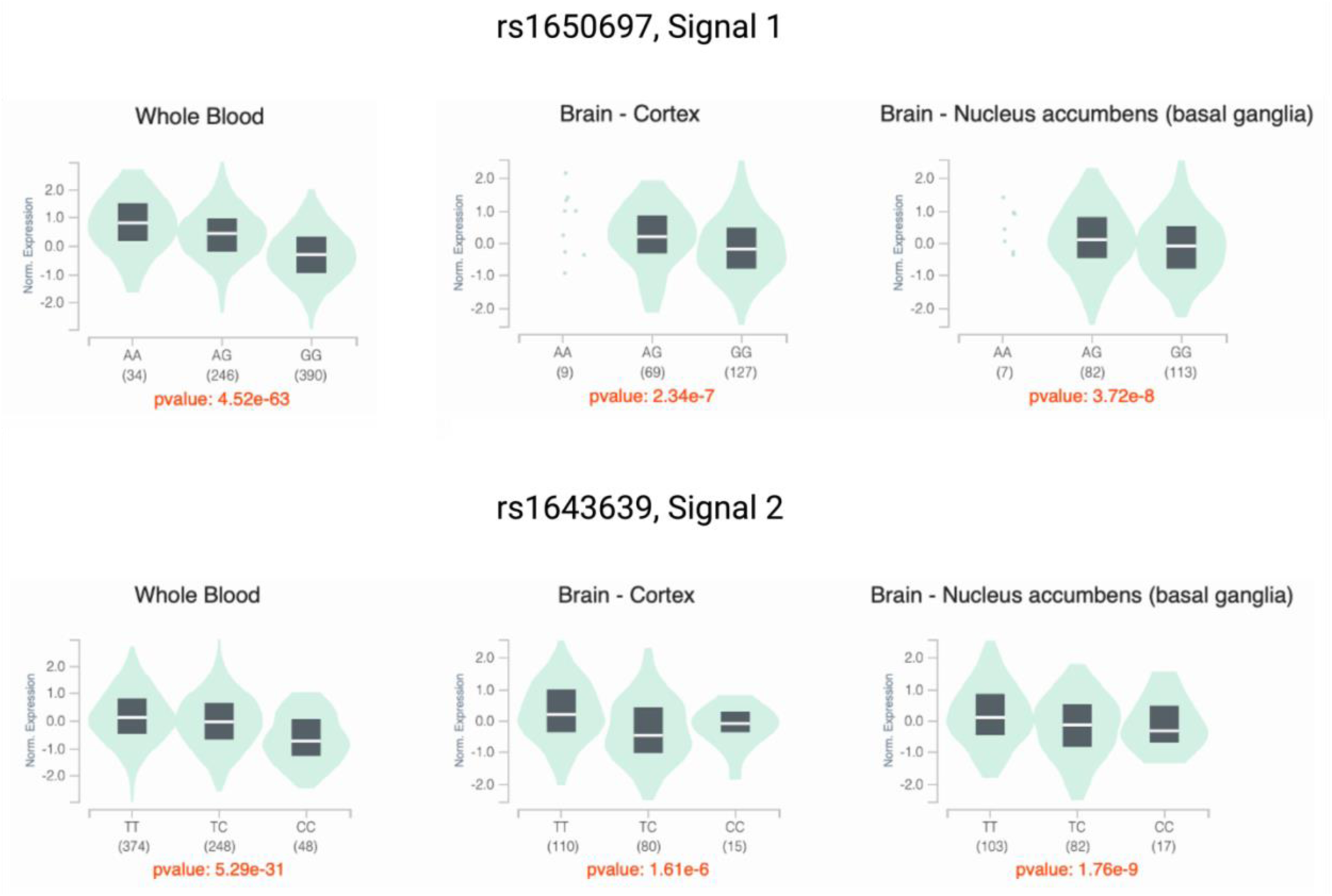
GTEx analyses of top signal 1 and signal 2 *MSH3* modifier variants. Graphs displaying correlations of MSH3 expression (y-axis) and rs1650697 and rs1643639 genotypes (x-axis). Numbers in brackets are the number of individuals with each genotype.

**Figure S10.**
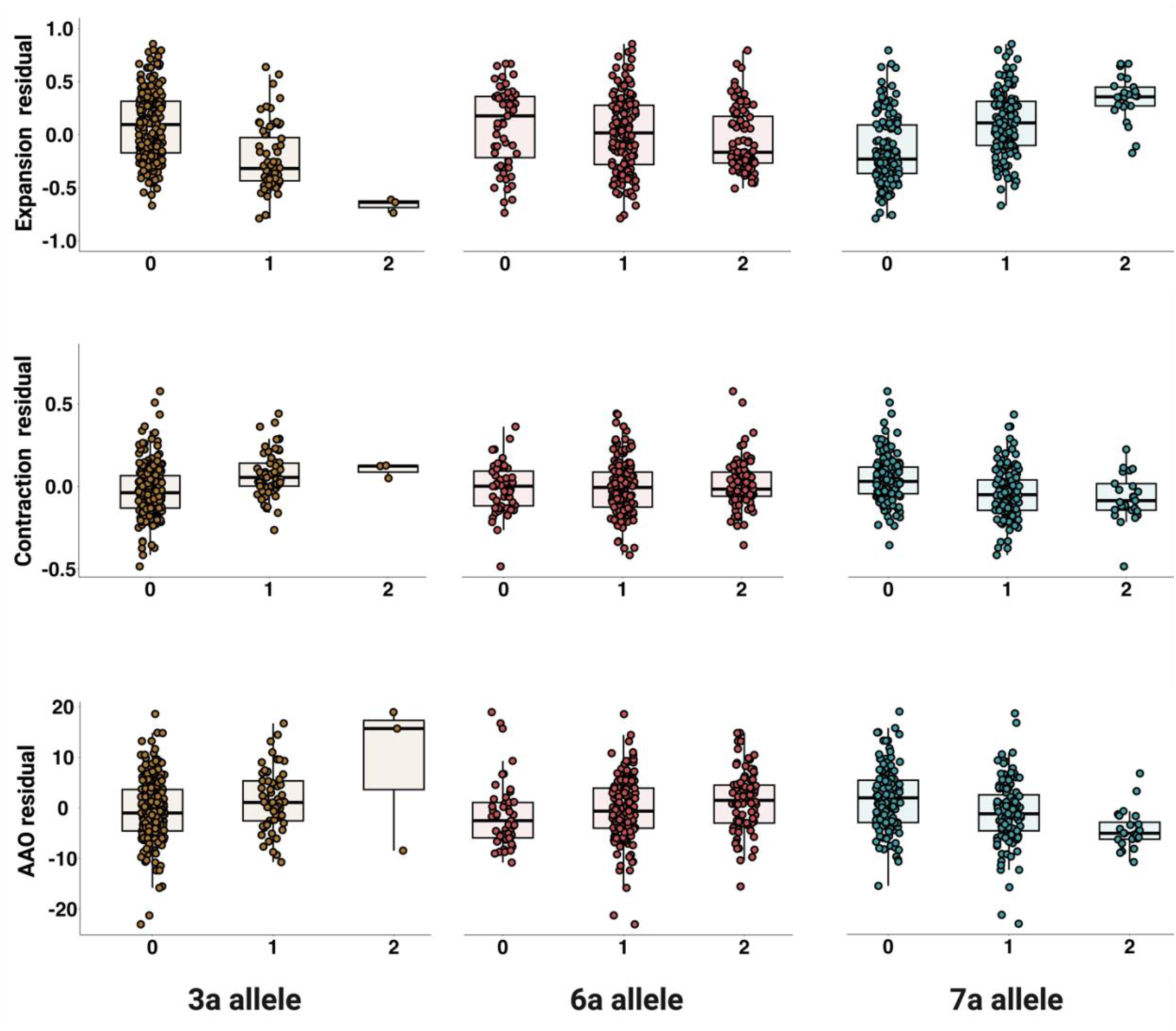
*MSH3* repeat variant allele dosage relationships with age at onset and somatic instability in blood. Expansion (top), contraction (middle) and AAO (bottom) residuals are plotted for individuals harboring 0, 1, or 2 alleles of the *MSH3* 3a, 6a or 7a repeat variant. See **Table S17** for statistical analyses.

**Figure S11.**
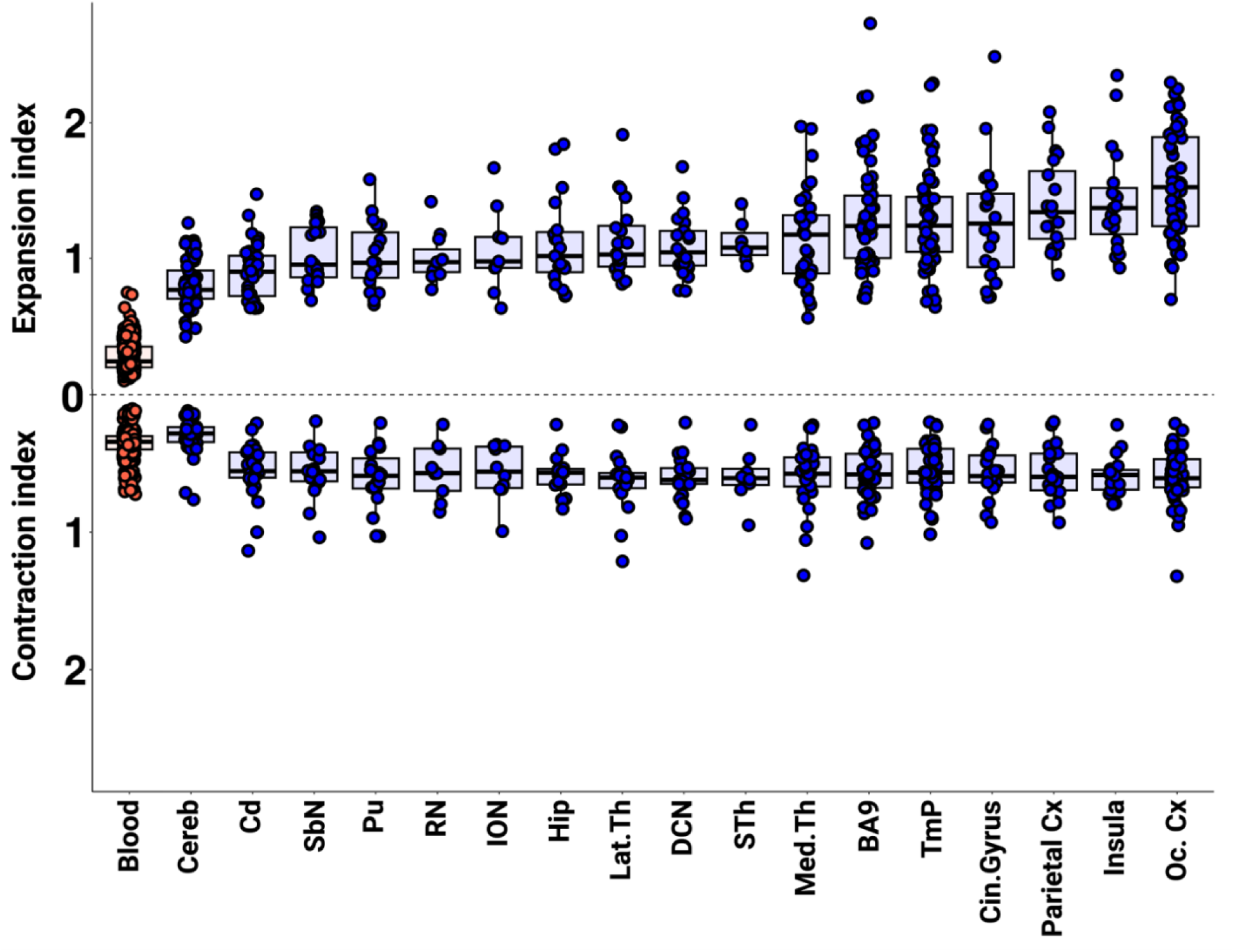
XDP CCCTCT repeat expansion and contraction in XDP brain tissues. Distribution of expansion indices (top panel) and contraction indices (bottom panel) ranked by median expansion index values. Note that for comparison with brain, the blood contraction index is calculated based on all the contraction peaks (see Materials and Methods). The expansion data parallel those previously reported^19^ but in a larger sample size to compare directly with the contraction data in the current study. Box-whisker plots show median ± interquartile range (IQR) and dots show values in individual patient samples. Cereb = cerebellum (n = 60), Cd = caudate (n = 33), SbN = substantia nigra (n = 19), ION = inferior olivary nucleus (n = 9), RN = red nucleus (n = 11), Med.Th = medial thalamus (n = 34), Hip = hippocampus (n = 19), Pu = putamen (n = 19), Lat.Th = lateral thalamus (n = 20), DCN = deep cerebellar nuclei (n = 21), STh = subthalamic nucleus (n = 7), Cin.Gyrus = cingulate gyrus (n = 20), BA9 = frontal cortex Brodmann area 9 (n = 58), Parietal Cx = parietal cortex (n = 20), Insula = insular cortex (n = 19), TmP = temporal pole (n = 57), Oc. Cx = occipital cortex (n = 60). Each dot represents an XDP individual.

**Figure S12.**
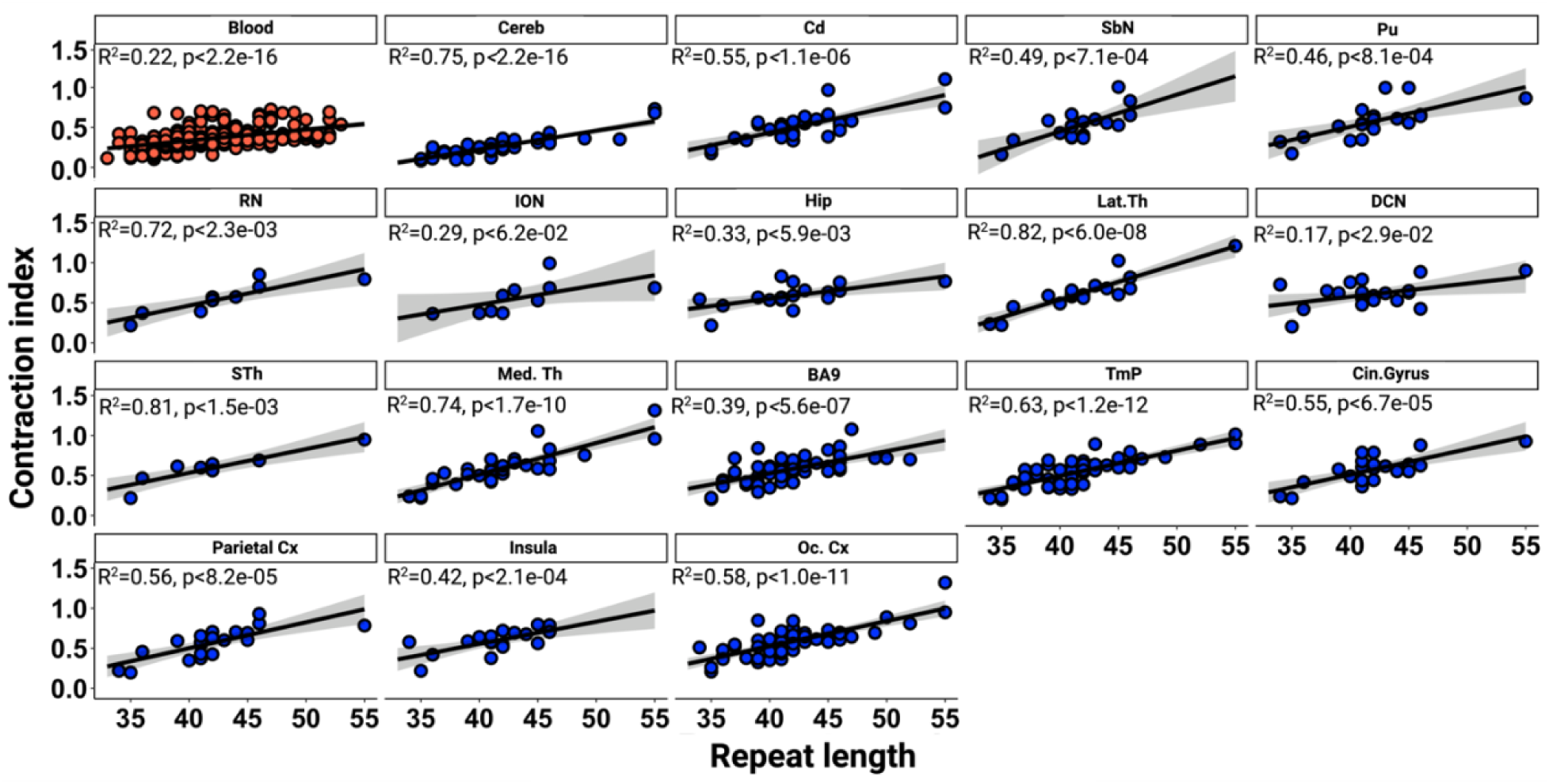
Linear regression analyses between CCCTCT inherited repeat length and contraction index in XDP brain tissues. All the regression equations, except for deep cerebellar nuclei, show significant association of contraction index (absolute value is plotted) with inherited repeat length. Grey shaded areas show 95% confidence interval.

**Figure S13.**
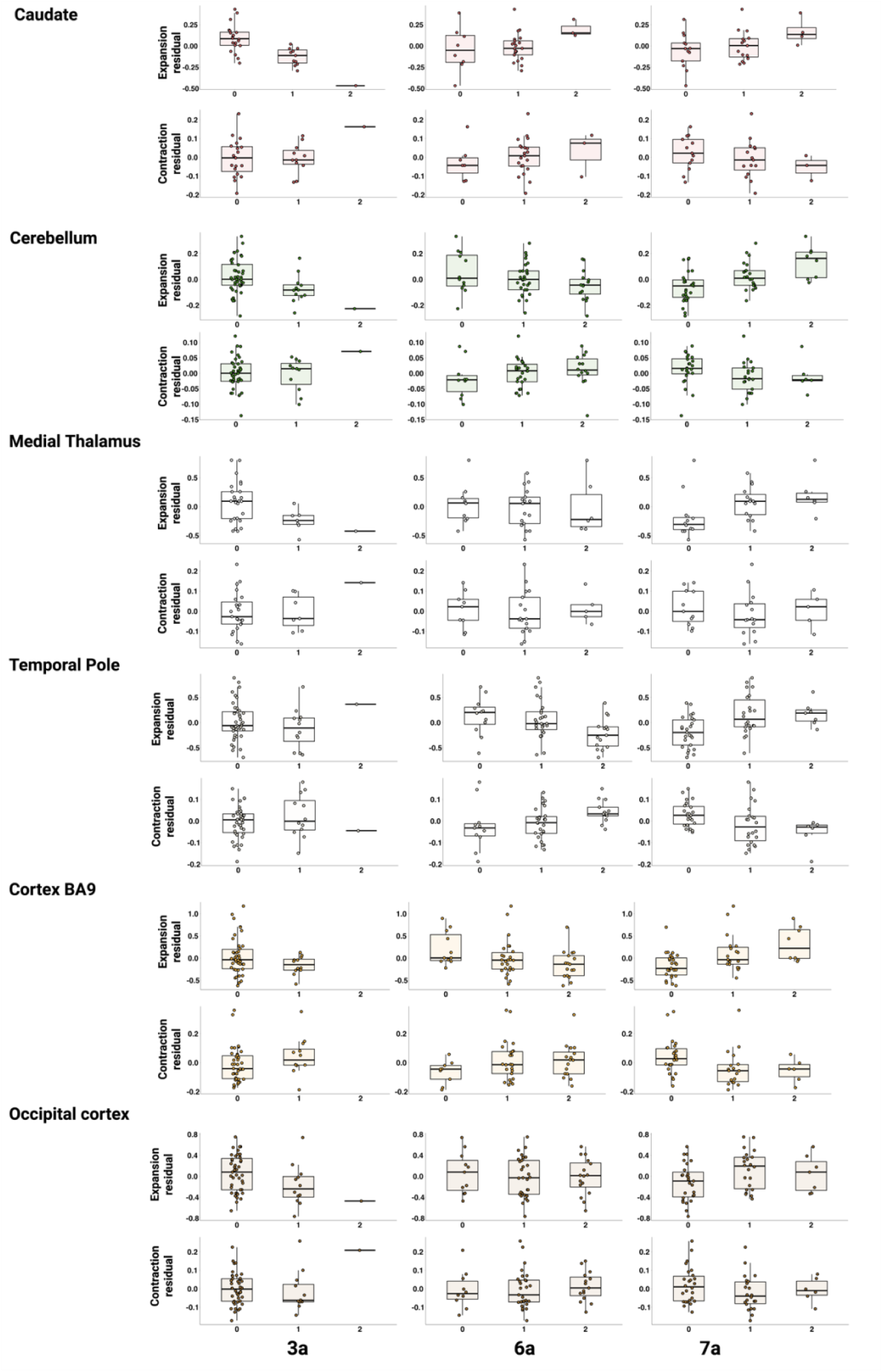
*MSH3* repeat variant allele dosage relationships with instability in brain. Expansion (top) and Contraction (bottom) residuals are plotted for individuals harboring 0, 1, or 2 alleles of the *MSH3* 3a, 6a or 7a repeat variant. See **Table S21** for statistical analyses.

